# Mapping the ecological networks of microbial communities

**DOI:** 10.1101/150649

**Authors:** Yandong Xiao, Marco Tulio Angulo, Jonathan Friedman, Matthew K. Waldor, Scott T. Weiss, Yang-Yu Liu

## Abstract

Microbes form complex and dynamic ecosystems that play key roles in the health of the animals and plants with which they are associated. Such ecosystems are often represented by a directed, signed and weighted ecological network, where nodes represent microbial taxa and edges represent ecological interactions. Inferring the underlying ecological networks of microbial communities is a necessary step towards understanding their assembly rules and predicting their dynamical response to external stimuli. However, current methods for inferring such networks require assuming a particular population dynamics model, which is typically not known a priori. Moreover, those methods require fitting longitudinal abundance data, which is not readily available, and often does not contain the variation that is necessary for reliable inference. To overcome these limitations, here we develop a new method to map the ecological networks of microbial communities using steady-state data. Our method can qualitatively infer the inter-taxa interaction types or signs (positive, negative or neutral) without assuming any particular population dynamics model. Additionally, when the population dynamics is assumed to follow the classic Generalized Lotka-Volterra model, our method can quantitatively infer the inter-taxa interaction strengths and intrinsic growth rates. We systematically validate our method using simulated data, and then apply it to four experimental datasets of microbial communities. Our method offers a novel framework to infer microbial interactions and reconstruct ecological networks, and represents a key step towards reliable modeling of complex, real-world microbial communities, such as the human gut microbiota.

## 1. Introduction

The microbial communities established in animals, plants, soils, oceans, and virtually every ecological niche on Earth perform vital functions for maintaining the health of the associated ecosystems^1-5^. Recently, our knowledge of the organismal composition and metabolic functions of diverse microbial communities has markedly increased, due to advances in DNA sequencing and metagenomics^6^. However, our understanding of the underlying ecological networks of these diverse microbial communities lagged behind^7^. Mapping the structure of those ecological networks and developing ecosystem-wide dynamic models will be important for a variety of applications^8^, from predicting the outcome of community alterations and the effects of perturbations^9^, to the engineering of complex microbial communities^7,10^. We emphasize that the ecological network discussed here is a directed, signed and weighted graph, where nodes represent microbial taxa and edges represent direct ecological interactions (e.g., parasitism, commensalism, mutualism, amensalism or competition) between different taxa. This is fundamentally different from the correlation-based association or co-occurrence network^7,11,12,13^, which is undirected and does not encode any causal relations or direct ecological interactions, and hence cannot be used to faithfully predict the dynamic behaviour of microbial communities.

To date, existing methods for inferring the ecological networks of microbial communities are based on temporal abundance data, i.e., the abundance time series of each taxon in the microbial community^14-19^. The success of those methods has been impaired by at least one of the following two fundamental limitations. *First*, those inference methods typically require the *a priori* choice of a parameterized population dynamics model for the microbial community. These choices are hard to justify, given that microbial taxa in the microbial community interact via a multitude of different mechanisms^7,20,21,22^, producing complex dynamics even at the scale of two taxa^23,24^. Any deviation of the chosen model from the “true” model of the microbial community can lead to systematic inference errors, regardless of the inference method that is used^19^. *Second*, a successful temporal-data based inference requires sufficiently informative time-series data^19,25^. For many host-associated microbial communities, such as the human gut microbiota, the available temporal data are often poorly informative. This is due to the fact that such microbial communities often display stability and resilience^26,27^, which leads to measurements containing largely their steady-state behavior. For microbial communities such as the human gut microbiota, trying to improve the informativeness of temporal data is challenging and even ethically questionable, as it requires applying drastic and frequent perturbations to the microbial community, with unknown effects on the host.

To circumvent the above fundamental limitations of inference methods based on temporal data, here we developed a new method based on *steady-state data*, which does not require any external perturbations. The basic idea is as follows. Briefly, if we assume that the net ecological impact of species on each other is context-independent, then comparing equilibria (i.e., steady-state samples) consisting of different subsets of species would allow us to infer the interaction types. For example, if one steady-state sample differs from another only by addition of one species X, and adding X brings down the absolute abundance of Y, then we can conclude X inhibits the growth of Y. This very simple idea can actually be extended to more complicated cases where steady-state samples differ from each other by more than one species. Indeed, we rigorously proved that, if we collect enough independent steady states of the microbial community, it is possible to infer the microbial interaction types (positive, negative and neutral interactions) and the structure of the ecological network, without requiring any population dynamics model. We further derived a rigorous criterion to check if the steady-state data from a microbial community is consistent with the Generalized Lotka-Volterra (GLV) model^15-19^, a classic population dynamics model for microbial communities in human bodies, soils and lakes. We finally proved that, if the microbial community follows the GLV dynamics, then the steady-state data can be used to accurately infer the model parameters, i.e., inter-taxa interaction strengths and intrinsic growth rates. We validated our inference method using simulated data generated from various classic population dynamics models. Then we applied it to real data collected from four different microbial communities.

## 2. Results

Microbes do not exist in isolation but form complex ecological networks^7^. The ecological network of a microbial community is encoded in its population dynamics, which can be described by a set of ordinary differential equations (ODEs):

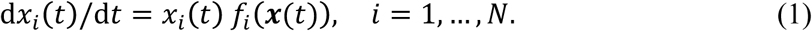

Here, *f*_*i*_(*x*(*t*)) ’s are some unspecified functions whose functional forms determine the structure of the underlying ecological network; 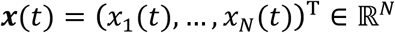 is an *N*-dimensional vector with *x*_*i*_(*t*) denoting the absolute abundance of the *i*-th taxon at time *t*. In this work, we don’t require ‘taxon’ to have a particular taxonomic ranking, as long as the resulting abundance profiles are distinct enough across all the collected samples. Indeed, we can group microbes by species, genus, family or just operational taxonomic units (OTUs).

Note that in the right-hand side of Eq. (1) we explicitly factor out *x*_*i*_ to emphasize that (i) without external perturbations those initially absent or later extinct taxa will never be present in the microbial community again as time goes by, which is a natural feature of population dynamics (in the absence of taxon invasion or migration); (ii) there is a trivial steady state where all taxa are absent; (iii) there are many non-trivial steady states with different taxa collections. We assume that the steady-state samples collected in a dataset 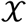 correspond to those non-trivial steady states ***x***^∗^ of Eq. (1), which satisfy 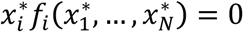, *i* = 1,…, *N*. For many host-associated microbial communities, e.g., the human gut microbiota, those cross-sectional samples collected from different individuals contain quite different collections of taxa (up to the taxonomic level of phylum binned from OTUs)^26^. We will show later that the number of independent steady-state samples is crucial for inferring the ecological network.

Mathematically, the intra- and inter-taxa ecological interactions (i.e., promotion, inhibition, or neutral) are encoded by the Jacobian matrix 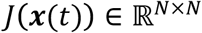 with matrix elements *J*_*ij*_(***x***(*t*)) = *∂f*_*i*_(***x***(*t*))/*∂x*_*j*_. The condition *J*_*ij*_(***x***(*t*)) > 0 (< 0 or = 0) means that taxon *j* promotes (inhibits or doesn’t affect) the growth of taxon *i*, respectively. The diagonal terms *J*_*ii*_(***x***(*t*)) represent intra-taxa interactions. Note that *J*_*ij*_(***x***(*t*)) might depend on the abundance of many other taxa beyond *i* and *j* (due to the so-called “higher-order” interactions^24,28-32^).

The structure of the ecological network is represented by the zero-pattern of *J*(***x***(*t*)). Under a very mild assumption that 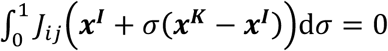 holds if and only if *J*_*ij*_ ≡ 0 (where ***x**^I^* and ***x**^k^* are two steady-state samples sharing taxon *i*), we find that the steady-state samples can be used to infer the zero-pattern of *J*(***x***(*t*)), i.e., the structure of the ecological network (see Supplementary Note 1.3 and 3 for details). Note that the network structure is interesting by itself and can be very useful in control theoretical analysis of microbial communities^33^. But in many cases, we are more interested in inferring the interaction types or strengths so that we can better predict the community’s response to perturbations.

The ecological interaction types are encoded in the sign-pattern of *J*(***x***(*t*)), denoted as sign(*J*(***x***(*t*))). To infer the interaction types, i.e., sign(*J*(***x***(*t*))), we make an explicit assumption that sign(*J*(***x***(*t*))) = const across all the observed steady-state samples. In other words, the nature of the ecological interactions between any two taxa does not vary across all the observed steady-state samples, though their interaction strengths might change. Note that the magnitude of *J*_*ij*_(***x***(*t*)) by definition may vary over different states, we just assume its sign remains invariant across all the observed samples/states. This assumption might be violated if those steady-state samples were collected from the microbial community under drastically different environmental conditions (e.g., nutrient availability^34^). In that case, inferring the interaction types becomes an ill-defined problem, since we have a “moving target” and different subsets of steady-state samples may offer totally different answers. Notably, as we will show later, the assumption is valid for many classic population dynamics models ^35-39^.

The assumption that sign(*J*(***x***(*t*))) = const can be falsified by analyzing steady-state samples. In Proposition 1 of Supplementary Note 1.4, we rigorously proved that if sign(*J*(***x***(*t*))) = const, then *true multi-stability* doesn’t exist. Equivalently, if a microbial community displays true multi-stability, then sign(*J*(***x***(*t*))) ≠ const. Here, a community of 0 taxa displays true multi-stability if there exists a subset of *M* (≤ *N*) taxa that has multiple different steady states, where all the *M* taxa have positive abundances and the other (*N* − *M*) taxa are absent. In practice, we can detect the presence of true multi-stability by examining the collected steady-state samples. If yes, then we know immediately that our assumption that sign(*J*(***x***(*t*))) = const is invalid and we should only infer the zero-pattern of *J*, i.e., the structure of the ecological network. If no, then at least our assumption is consistent with the collected steady-state samples, and we can use our method to infer sign(*J*(***x***(*t*))), i.e., the ecological interaction types. In short, by introducing a criterion to falsify our assumption, we significantly enhance the applicability of our method (see Supplementary Note 1.4 and Remark 6 for more detailed discussions).

### Inferring interaction types

The assumption that sign(*J*(***x***(*t*))) = const enables us to mathematically prove that sign(*J*(***x***(*t*))) satisfies a strong constraint (Theorem 2 in Supplementary Note 1.4). By collecting enough independent steady-state samples, we can solve for the sign-pattern of *J*(***x***) and hence map the structure of the ecological network (Remarks 4 and 5 in Supplementary Note 1.4).

The basic idea is as follows. Let *J*_*i*_ be the set of all steady-state samples sharing taxon *i*. Then, for any two of those samples ***x***^*I*^ and ***x***^*K*^, where the superscripts *I,K* ∈ *J*_*i*_ denote the collections of present taxa in those samples, we can prove that the sign-pattern of the *i*-th row of Jacobian matrix, denoted as a ternary vector ***s***_*i*_ ∈ {−, 0, +}^*N*^, is *orthogonal* to (***x***^*I*^ − ***x**^K^*) (Eq. (S3) in Supplementary Note 1.1). In other words, we can always find a real-valued vector 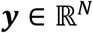, which has the same sign-pattern as ***s***_*i*_ and satisfies **y**^T^ · (***x**^I^* − ***x**^K^*) = 0. If we compute the sign-patterns of all vectors orthogonal to (***x**^I^* − ***x**^K^*) for all *I, K* ∈ *J*_*i*_, then ***s***_*i*_ must belong to the intersections of those sign-patterns, denoted as 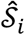. In fact, as long as the number Ω of steady-state samples in 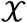 is above certain threshold Ω*, then 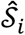 will contain only three sign-patterns {−***a*, 0, *a***} (Remark 5 in Supplementary Note 1.4). To decide which of these three remaining sign-patterns is the true one, we just need to know the sign of only one non-zero interaction. If such prior knowledge is unavailable, one can at least make a reasonable assumption that *s*_*ii*_ = ‘−’, i.e., the intra-taxa interaction *J*_*ii*_ is negative (which is often required for community stability). When 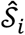 has more than three sign-patterns, we proved that the steady-state data is not informative enough in the sense that all sign-patterns in 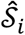 are consistent with the data available in 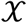 (Remark 5 in Supplementary Note 1.4). This situation is not a limitation of any inference algorithm but of the data itself. To uniquely determine the sign-pattern in such a situation, one has to either collect more samples (thus increasing the informativeness of 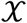) or use *a priori* knowledge of non-zero interactions.

We illustrate the application of the above method to small microbial communities with unspecified population dynamics (Fig. 1). For the two-taxa community (Fig. 1a), there are three possible types of equilibria, i.e., {***x***^{1}^, ***x***^{2}^, ***x***^{1,2}^}, depicted as colored pie charts in Fig. 1b. In order to infer ***s***_1_ = (sign(*J*_11_), sign(*J*_12_)), we compute a straight line (shown in green in Fig. 1b) that is orthogonal to the vector (***x***^{1,2}^ − ***x***^{1}^) and passes through the origin. The regions (including the origin and two quadrants) crossed by this green line provide the set of possible sign-patterns 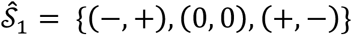 that ***s***_1_ may belong to. A *priori* knowing that *J*_11_ < 0, our method correctly concludes that ***s***_1_ = (−, +). Note that *J*_12_ > 0 is consistent with the observation that with the presence of taxon 2, the steady-state abundance of taxon 1 increases (Fig. 1b), i.e., taxon 2 promotes the growth of taxon 1. We can apply the same method to infer the sign-pattern of ***s***_2_ = (−, −).

**Figure 1.**
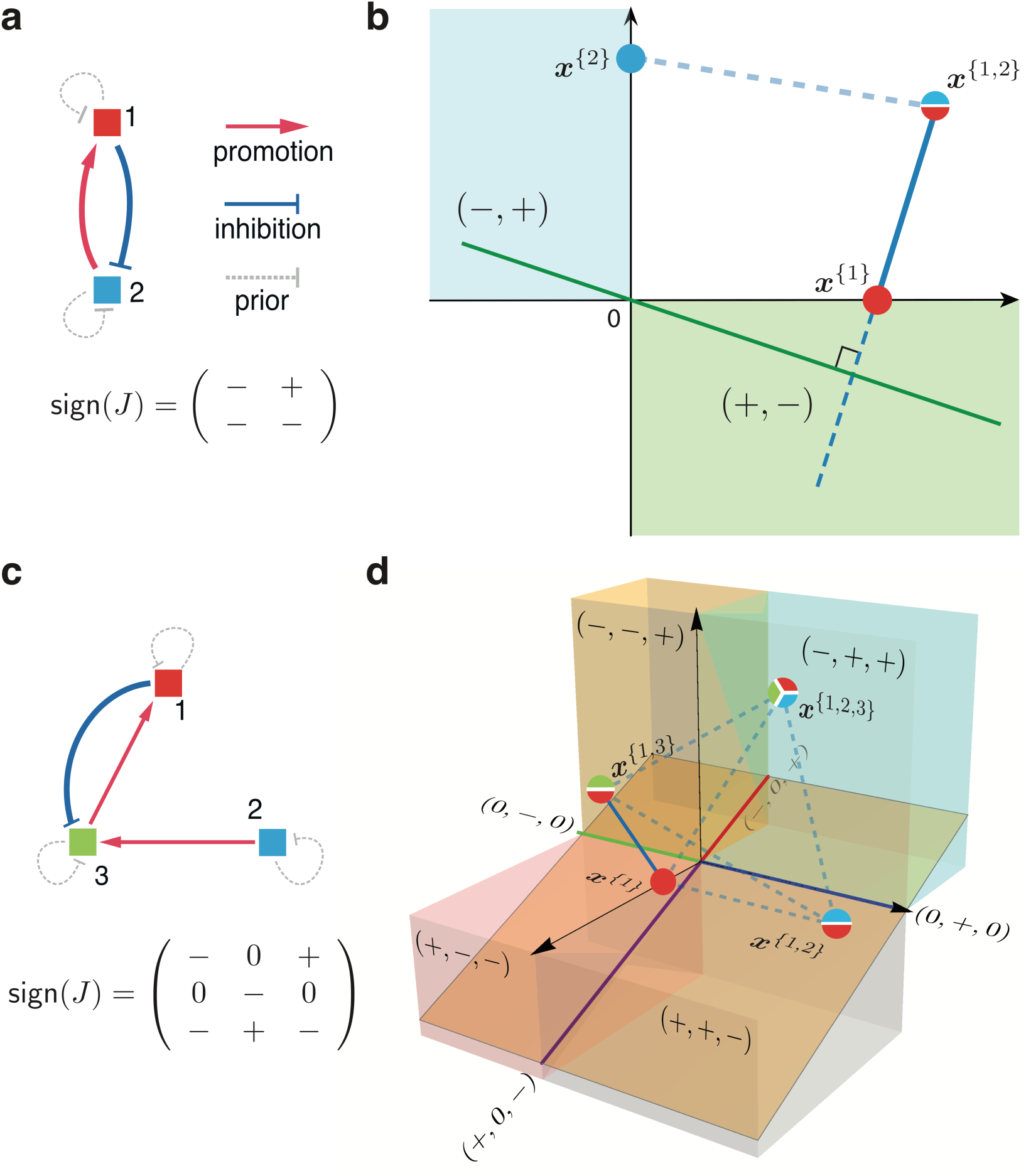
Inferring ecological interaction types for a small microbial community. The interaction types are coded as the sign-pattern of the Jacobian matrix. **a.** For a microbial community of 2 taxa, its ecological network and the sign-pattern of the corresponding Jacobian matrix are shown here. **b.** There are three possible steady-state samples (shown as colored pie charts), and two of them ***x***^{1,2}^, ***x***^{1}^ share taxon 1. We can calculate the green line that passes through the origin and is perpendicular to the vector (***x***^{1,2}^ − ***x***^{1}^) (shown as a blue line segment). This green line crosses the origin, and two other orthants (shown in light cyan and green), offering a set of possible sign-patterns: (0,0), (−,−) and (+, −), for which **s**_1_ = (sign(*J*_11_), sign(*J*_12_)) may belong to. Provided that *J*_11_ < 0, we conclude that ***s***_1_ = (−,+). **c.** For a microbial community of 3 taxa, its ecological network and the sign-pattern of the corresponding Jacobian matrix are shown here. **d**. There are seven possible steady-state samples, and we plot four of them that share taxon 1. Consider a line segment (***x***^{1,3}^ − ***x***^{1}^) (solid blue). We calculate the orange plane that passes through the origin and is perpendicular to this solid blue line. This orange plane crosses 9 regions: the origin and the other 8 regions (denoted in different color cubes, color lines), offering 9 possible sign-patterns for **s**_1_. We can consider another line segment that connects two steady-state samples sharing taxon 1, say, ***x***^{1,3}^ and ***x**^{1,2,3}^*, and repeat the above procedure. We do this for all the sample pairs (dashed blue lines), record the regions crossed by the corresponding orthogonal planes. Finally, the intersection of the regions crossed by all those orthogonal hyperplanes yields a minimum set of sign-patterns 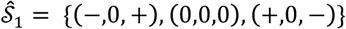 that ***s***_1_ may belong to. If we know that *J*_11_ < 0, then we can uniquely determine ***s***_1_ = (−,0, +).

For the three-taxa community (Fig. 1c), there are seven possible types of equilibria, i.e., {***x***^{1}^, ***x***^{2}^, ***x***^{3}^, ***x***^{1,2}^, ***x***^{1,3}^, ***x***^{2,3}^, ***x***^{1,2,3}^}. Four of them share taxon 1 (see colored pie charts in Fig. 1d). Six line segments connect the 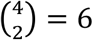 sample pairs, and represent vectors of the form (***x***^*I*^ − ***x***^*K*^), *I, K* ∈ *J*_1_ ={{1},{1,2},{1,3},{1,2,3}}. Considering a particular line segment (***x***^{1,3}^ − ***x***^{1}^), i.e., the solid blue line in Fig. 1d, we compute a plane (shown in orange in Fig. 1d) that is orthogonal to it and passes through the origin. The regions (including the origin and eight orthants) crossed by this orange plane provide a set of possible sign-patterns that ***s***_1_ may belong to (see Fig. 1d). We repeat the same procedure for all other vectors (***x**^I^* − ***x**^K^*), *I, K* ∈ *J*_1_, and compute the intersection of all the possible sign-patterns, finally yielding the minimum set 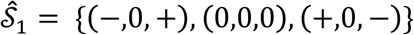 to which ***s***_1_ may belong to. If the sign of one non-zero interaction is known (*J*_11_ < 0 for this example), our method correctly infers the true sign-pattern ***s***_1_ = (−,0,+). Repeating this process for samples sharing taxon 2 (or 3) will enable us to infer the sign-pattern ***s***_2_ (or ***s***_3_), respectively.

It is straightforward to generalize the above method to a microbial community of *N* taxa (see Supplementary Note 2.1 for details). But this brute-force method requires us to calculate all the sign-pattern candidates first, and then calculate their intersection to determine the minimum set 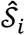 that ***s***_*i*_ will belong to. Since the solution space of sign-patterns is of size 3^*N*^, the time complexity of this brute force method is exponential with *N*, making it impractical for a microbial community with *N* > 10 taxa (Supplementary Note 2.2). To resolve this issue, we developed a heuristic algorithm that pre-calculates many intersection lines of (*N* − 1) non-parallel hyperplanes that pass through the origin and are orthogonal to (***x**^I^* − ***x**^K^*), *I*, *K* ∈ *j*_*i*_. Based on these pre-calculated intersection lines, the algorithm determines 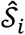 using the most probable intersection line. The solution space of this heuristic algorithm is determined by the user-defined number of pre-calculated interaction lines (denoted as Ψ). Hence this algorithm naturally avoids searching the exponentially large solution space (see Supplementary Note 2.3 for details). Later on, we will show that this heuristic algorithm can indeed infer the interaction types with high accuracy.

In reality, due to measurement noise and/or transient behavior of the microbial community, the abundance profiles of the collected samples may not exactly represent steady states of the microbial community. Hence for certain *J*_*ij*_ ’s their inferred signs might be wrong. Using simulated data, we will show later that for considerable noise level the inference accuracy is still reasonably high.

### Inferring interaction strengths

To quantitatively infer the inter-taxa interaction strengths, it is necessary to choose *a priori* a parameterized dynamic model for the microbial community. The classical GLV model can be obtained from Eq. (1) by choosing

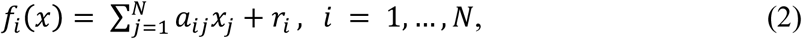

where 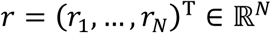 is the intrinsic growth rate vector and 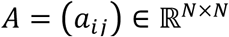 is the interaction matrix characterizing the intra- and inter-taxa interactions.

From Eq. (2) we can easily calculate the Jacobian matrix *J*, which is nothing but the interaction matrix *A* itself. This also reflects the fact that the value of *a*_*ij*_ quantifies the interaction strength of taxon *j* on taxon *i*. The GLV model considerably simplifies the inference of the ecological network, because we can prove that ***a***_*i*_ · (***x**^1^* − ***x**^K^*) = 0, for all *I, K* ∈ *J*_*i*_, where ***a***_*i*_ ≡ (*a*_*i*1_,…, *a*_*iN*_) represents the *i*-th row of *A* matrix (Supplementary Note 5.2). In other words, all steady-state samples containing the *i*-th taxon will align exactly onto a hyperplane, whose orthogonal vector is parallel to the vector ***a***_*i*_ that we aim to infer (Fig. 2a, Theorem 3 of Supplementary Note 5.1). Thus, for the GLV model, the inference from steady-state data reduces to finding an (*N* − 1)-dimensional hyperplane that “best fits” the steady-state sample points {***x**^I^*|*I* ∈ *J*_*i*_} in the *N*-dimensional state space. In order to exactly infer ***a***_*i*_, it is necessary to know the value of at least one non-zero element in ***a***_*i*_, say, ***a***_*ii*_. Otherwise, we can just determine the *relative* interaction strengths by expressing *a*_*ij*_ in terms of *a*_*ii*_. Once we obtain ***a***_*i*_, the intrinsic growth rate *r*_*i*_ of the *i*-th taxon can be calculated by averaging (−***a***_*i*_ · ***x**^I^*) over all *I ∈ J*_*i*_, i.e., all the steady-state samples containing taxon *i*.

**Figure 2.**
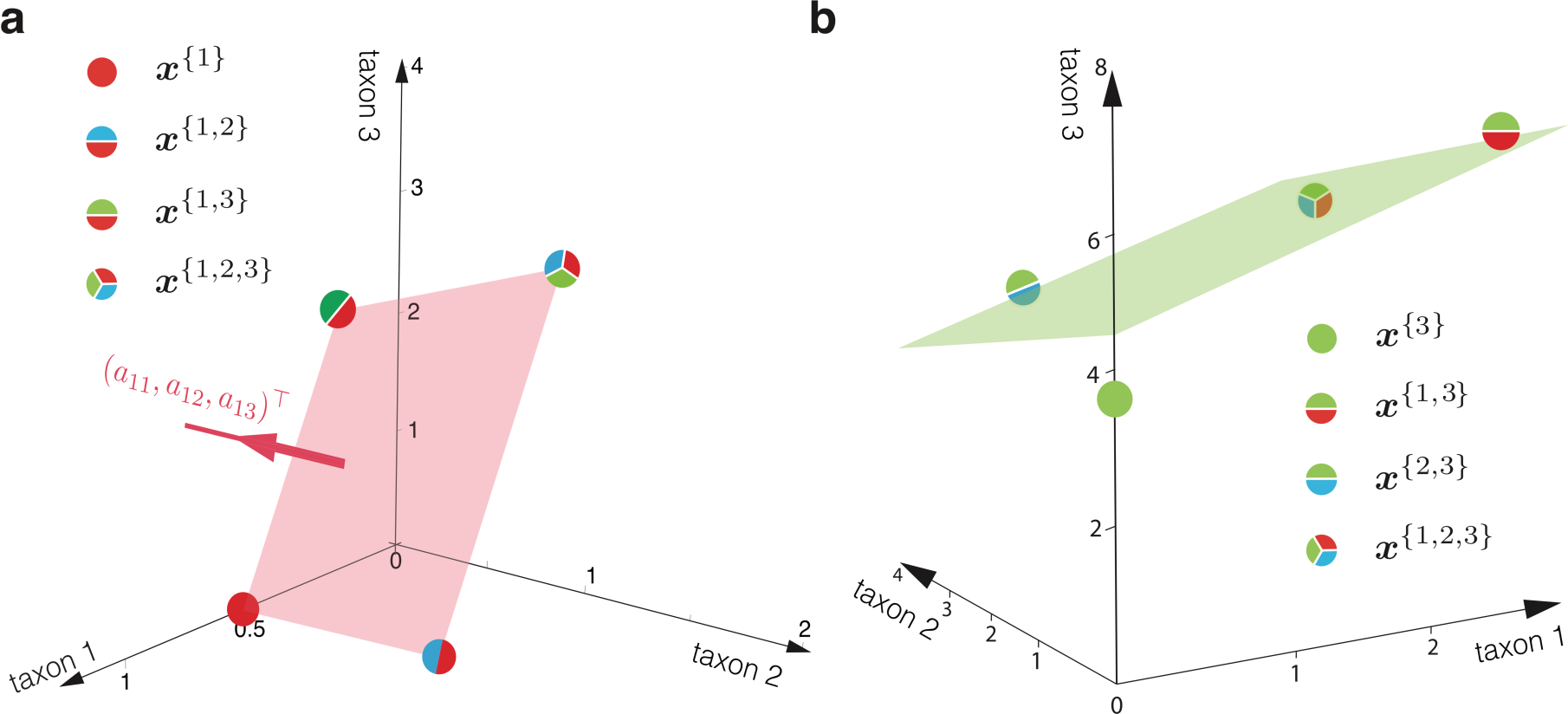
Consistency check of the GLV model and the observed steady-state samples. For a microbial community following exactly the GLV dynamics, all its steady-state samples sharing one common taxon will align onto a hyperplane in the state space. **a.** Here we consider a microbial community of three taxa. There are four steady-state samples {***x***^{1}^, ***x***^{1,2}^,***x***^{1,3}^,***x***^{1,2,3}^} that share common taxon 1. Those four steady-state samples represent four points in the state space, and they align onto a plane (light red). The normal vector of this plane is parallel to the first row ***a***_1_ of the interaction matrix j in the GLV model. Given any one of non-zero entries in *a*_1_, we can determine the exact values of all other entries. Otherwise, we can always express the intertaxa interaction strengths ***a***_*ij*_ (*j* ≠ *i*) as a function of the intra-taxa interaction strength ***a***_*ii*_. **b.** Here we again consider a microbial community of three taxa. Taxon-1 and taxon-2 follow the GLV dynamics, but taxon-3 doesn’t. Then those steady-state samples that share common taxon-3 will not align onto a plane anymore. Here we show the best fitted plane (in green) by minimizing the distance between this plane and the four steady states, with the coefficient of determination *R*^2^ = 0.77.

In case the samples are not collected exactly at steady states of the microbial community or there is noise in abundance measurements, those samples containing taxon *i* will not exactly align onto a hyperplane. A naive solution is to find a hyperplane that minimizes its distance to those noisy samples. But this solution is prone to induce false positive errors and will yield non-sparse solutions (corresponding to very dense ecological networks). This issue can be partly alleviated by introducing a Lasso regularization^40^, implicitly assuming that the interaction matrix *A* in the GLV model is sparse. However, the classical Lasso regularization may induce a high false discovery rate (FDR), meaning that many zero interactions are inferred as non-zeros ones. To overcome this drawback, we applied the Knockoff filter^41^ procedure, allowing us to control the FDR below a desired user-defined level *q* > 0 (see Supplementary Note 5.3 for details).

The observation that for the GLV model all noiseless steady-state samples containing the *i*-th taxon align exactly onto a hyperplane can also be used to characterize how much the dynamics of the *i*-th taxon in a real microbial community deviates from the GLV model. This deviation can be quantified by the coefficient of determination (denoted by *R*^2^) of the multiple linear regression when fitting the hyperplane using the steady-state samples (Fig. 2b). If *R*^2^ is close to 1 (the samples indeed align to a hyperplane), we conclude that the dynamics of the microbial community is consistent with the GLV model, and hence the inferred interaction strengths and intrinsic growth rates are reasonable. Otherwise, we should only aim to qualitatively infer the ecological interaction types that do not require specifying any population dynamics.

### Validation on simulated data

#### Interaction types

To validate the efficacy of our method in inferring ecological interaction types, we numerically calculated the steady states of a small microbial community with *N* = 8 taxa, using four different population dynamics models^35-39^: Generalized Lotka-Volterra (GLV), Holling Type II (Holling II), DeAngelis-Beddington (DB) and Crowley-Martin (CM) models (see Supplementary Note 4 for details). Note that all these models satisfy the requirement that the sign-pattern of the Jacobian matrix is time-invariant. To infer the ecological interaction types among the 8 taxa, we employed both the brute-force algorithm (with solution space ~ 3^8^ = 6,561) and the heuristic algorithm (with solution space given by the number of the pre-calculated intersections chosen as Ψ = 5*N* = 40).

In the noiseless case, we find that when the number of steady-state samples satisfies Ω > 3*N*, the heuristic algorithm outperformed the brute-force algorithm for datasets generated from all the four different population dynamics models (Fig. 3a). This result is partly due to the fact that the former requires much fewer samples than the latter to reach high accuracy (the percentage of correctly inferred interaction types). However, when the sample size Ω is small (< 3*N*), the heuristic algorithm completely fails while the brute-force algorithm still works to some extent.

**Figure 3.**
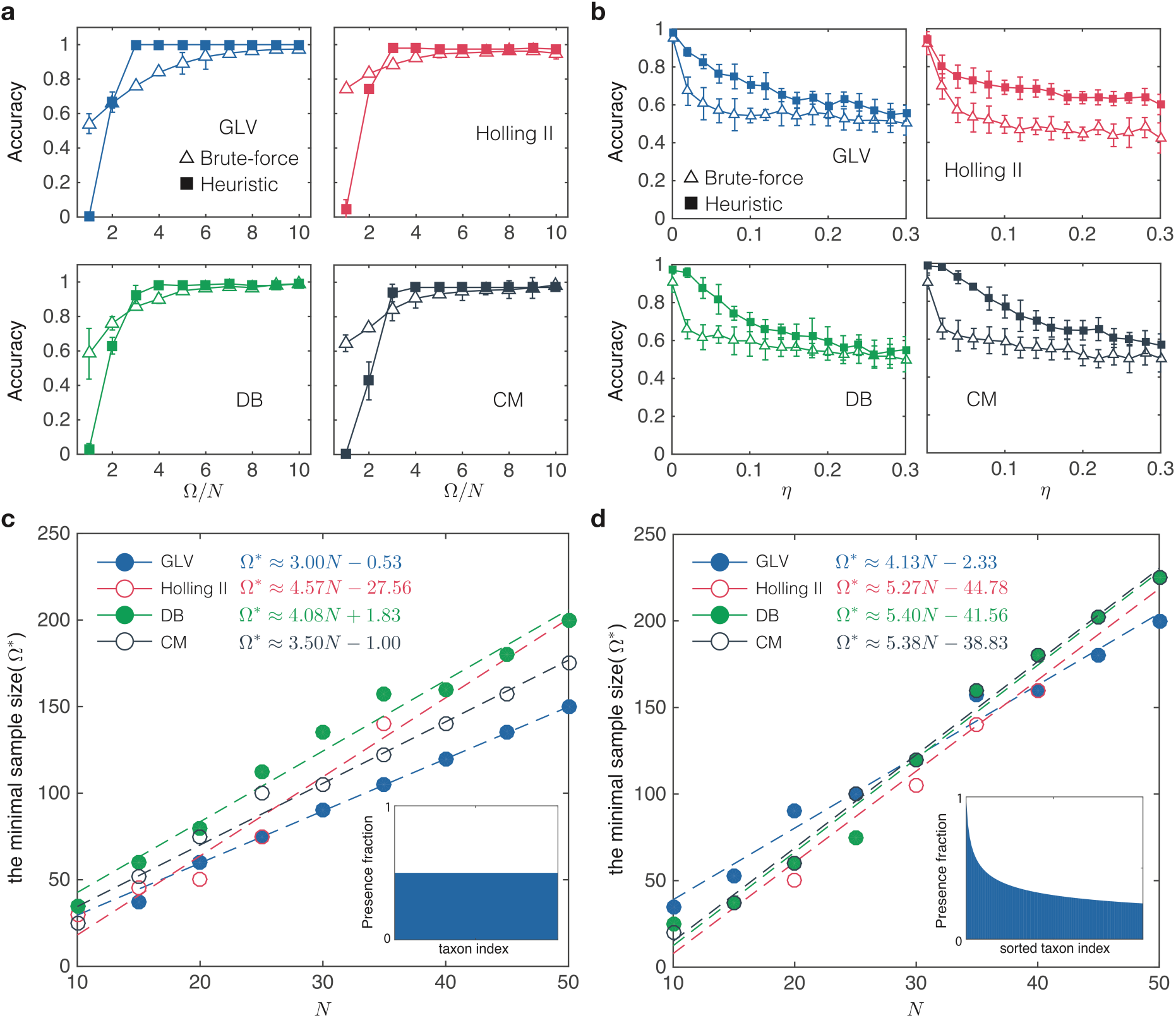
Validation of our method in inferring interaction types using simulated data. a-b: Consider a small microbial community of *N* = 8 taxa. We generate steady-state samples using four different population dynamics models: Generalized Lotka-Volterra (GLV), Holling Type II (Holling II), DeAngelis-Beddington (DB) and Crowley-Martin (CM). We compare the performance of the brute-force algorithm (with solution space ~3^8^ = 6,561) and the heuristic algorithm (with solution space ~ Ψ = 5*N* = 40). **a.** In the noiseless case, we plot the inference accuracy (defined as the percentage of correctly inferred signs in the Jacobian matrix) as a function of sample size Ω. **b.** In the presence of noise, we plot the inference accuracy as a function of the noise level *η*. Here the sample size is fixed: Ω = 5*N* = 40. The error bar represents standard deviation for 10 different realizations. **c-d.** We calculate the minimal sample size Ω* required for the heuristic algorithm to achieve high accuracy (100% for GLV, 95% for Holling II, DB and CM) at different system sizes. We consider two different taxa presence patterns: uniform and heterogeneous (see insets). Here the simulated data is generated in the noiseless case and we chose Ψ = 10*N*. a-d: The underlying ecological network is generated from a directed random graph model with connectivity 0.4 (i.e., with probability 0.4 there will be a directed edge between any two taxa).

We then fix Ω = 5*N*, and compare the performance of the brute-force and heuristic algorithms in the presence of noise (Fig. 3b). We add artificial noise to each non-zero entry 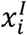 of a steady-state sample ***x**^1^* by replacing 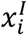 with 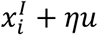, where 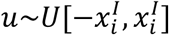 is a random number uniformly distributed in the interval 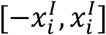 and *η* ≥ 0 quantifies the noise level. We again find that the heuristic algorithm works better than the brute-force algorithm for datasets generated from all the four different population dynamics models.

The above encouraging results on the heuristic algorithm prompt us to systematically study the key factor to obtain an accurate inference, i.e., the minimal sample size Ω* (Fig. 3c, d). Note that for a microbial community of *N* taxa, if we assume that for any subset of the *N* taxa there is only one stable steady state such that all the corresponding taxa have non-zero abundance, then there are at most Ω_max_ = (2^*N*^ − 1) possible steady-state samples (Of course, not all of them will be ecologically feasible. For example, certain pair of taxa will never coexist.) In general, it is unnecessary to collect all possible steady-state samples to obtain a highly accurate inference result. Instead, we can rely on a subset of them. To demonstrate this, we numerically calculated the minimal sample size Ω* we need to achieve a highly accurate inference of interaction types. We considered two different taxa presence patterns: (1) *uniform:* all taxa have equal probability of being present in the steady-state samples (inset of Fig. 3c); and (2) *heterogeneous:* a few taxa have higher presence probability than others, reminiscent of human gut microbiome samples^26^ (inset of Fig. 3d). We found that for the steady-state data generated from all the four population dynamics models, Ω* always scales linearly with *N* in both taxa presence patterns, and the uniform taxa presence pattern requires much fewer samples (Fig. 3c,d).

Note that as *N* grows, the total possible steady-state samples Ω_max_ increases exponentially, while the minimal sample size Ω* we need for high inference accuracy increase linearly. Hence, interestingly, we have Ω*/Ω_max_ → 0 as *N* increases. This suggests that as the number of taxa increases, the proportion of samples needed for accurate inference actually decreases. This is a rather counter-intuitive result because, instead of a “curse of dimensionality”, it suggests that a “blessing of dimensionality” exists when using the heuristic algorithm to infer interaction types for microbial communities with a large number of taxa.

#### Interaction strengths

To validate our method in quantitatively inferring inter-taxa interaction strengths, we numerically calculated steady states for a microbial community of *N* = 50 taxa, using the GLV model with ***a***_*ii*_ = − 1 for all taxa.

In the noiseless case, if during the inference we know exactly ***a***_*ii*_ = −1 for all taxa, then we can perfectly infer the inter-taxa interaction strengths ***a***_*ij*_’s and the intrinsic growth rates *r*_*i*_’s (see Fig. 4a). To study the minimal sample size Ω* required for perfect inference in the noiseless case, we again consider two different taxa presence patterns: (1) uniform; (2) heterogeneous. We find that for both taxa presence patterns Ω* scales linearly with *N*, though the uniform taxa presence pattern requires much fewer samples (Fig. 4b).

**Figure 4.**
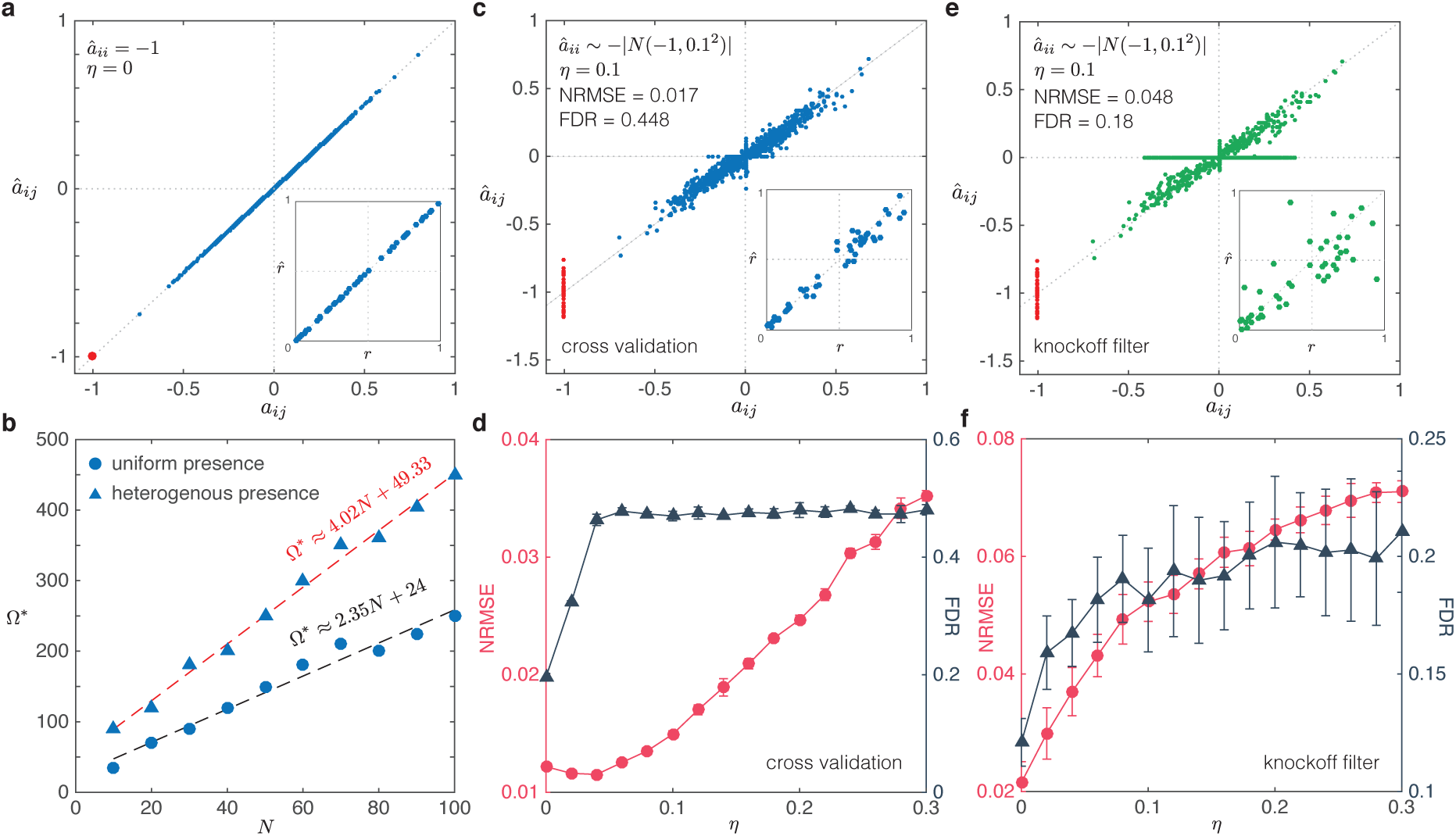
Validation of our method in inferring interaction strengths using simulated data. Here we simulate steady-state samples using the GLV model with *N* taxa and intra-taxa interaction strength *a*_*ii*_ = −1 for each taxon. The underlying ecological network is generated from a directed random graph model with connectivity 0.4. The noise is added to steady-state samples as follows: 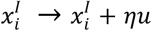, where the random number *u* follows a uniform distribution 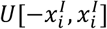, and *η* is the noise level. **a-b,** Consider the ideal case: (1) noiseless *η* = 0; and (2) we know exactly ***a***_*ii*_ = −1. **a.** We can perfectly infer *a*_*ij*_’s and *r*_*i*_’s. **b.** The minimal sample size Ω* required to correctly infer the interaction strengths scales linearly with the system size *N*. Here we consider two different taxa presence pattern: uniform and heterogeneous. **c-f**, In the presence of noise, and during the we just assume that the intra-taxa interaction strengths 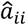 follows a half-normal distribution. **c**, Using the Lasso regularization induces high false discovery rate (FDR) ~ 0.448 at *η* = 0.1. **d.** Using classical Lasso with cross validation, both NRMSE and FDR increase with increasing *η*. **e.** For the same dataset used in (c), we use the knockoff filter to control the FDR below a certain level *q =* 0.2. **f.** With increasing noise level t, FDR can still be successfully controlled below *q* = 0.2 by applying the knockoff filter. In subfigures a,c-f, we have *N* = 50. The error bar represents standard deviation for 10 different realizations.

In the presence of noise, and if we don’t know the exact values of *a*_*ii*_’s, but just assume they follow a half-normal distribution 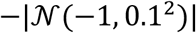, we can still infer *a*_*ij*_’s and *r*_*i*_’s with reasonable accuracy (with the normalized root-mean-square error NRMSE < 0.08), for noise level *η* < 0.3 (Fig. 4c-f). However, we point out that the classical Lasso regularization could induce many false positive, and the false discovery rate (FDR) reaches 0.448 at noise level *η* = 0.1, indicating that almost half of inferred non-zero interactions are actually zero (Fig. 4c). Indeed, even with a noise level *η* = 0.04, the classical Lasso already yields FDR~0.45, staying there for higher *η* (Fig. 4d).

In many cases, we are more concerned about low FDR than high false negative rates, because the topology of an inferred ecological network with even many missing links can still be very useful in the study of its dynamical and control properties^42^. To control FDR below a certain desired level *q* = 0.2, we applied the Knockoff filter^41^ (Fig. 4e), finding that though it will introduce more false negatives (see the horizontal bar in Fig. 4e), it can control the FDR below 0.2 for a wide range of noise level (Fig. 4f).

We also found that applying this GLV inference method to samples obtained from a microbial community with non-GLV dynamics leads to significant inference errors even in the absence of noise (Supplementary Fig. 9).

### Application to experimental data

#### A synthetic soil microbial community of eight bacterial species^43^

This dataset consists of steady states of a total of 101 different species combinations: all 8 solos, 28 duos, 56 trios, all 8 septets, and 1 octet (see Supplementary Note 6.1 for details). For those steady-state samples that started from the same species collection but with different initial conditions, we average over their final steady states to get a representative steady state for this particular species combination.

In the experiments, it was found that several species grew to a higher density in the presence of an additional species than in monoculture. The impact of each additional species (competitor) *j* on each focal species *i* can be quantified by the so-called ***relative yield**,* defined as: 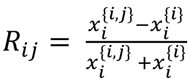, which represents a proxy of the ground truth of the interaction strength that species *j* impacts species *i*. A negative relative yield indicates growth hindrance of species *j* on *i*, whereas positive values indicated facilitation (Fig. 5a). Though quantifying the relative yield is conceptually easy and implementable for certain small microbial communities (see Supplementary Note 7 for details), for many host-associated microbial communities with many taxa, such as the human gut microbiota, measuring these one- and two-species samples is simply impossible. This actually motivates the inference method we developed here.

**Figure 5.**
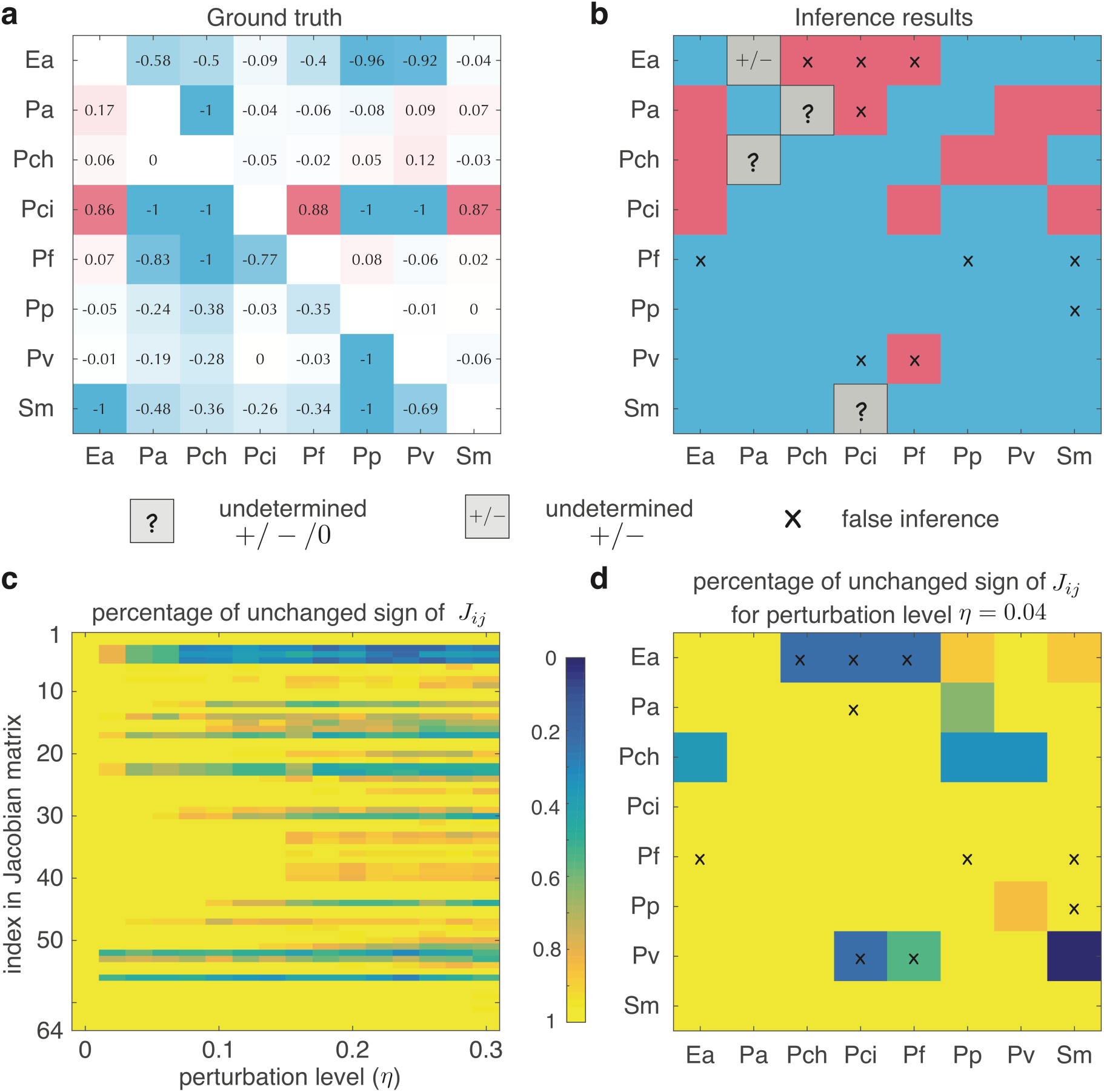
Inferring interaction types of a synthetic soil microbial community. The steady-state samples were experimentally collected from a synthetic soil microbial community of eight bacterial species. Those steady-state samples involve 101 different species combinations: all 8 solos, 28 duos, 56 trios, all 8 septets, and 1 octet. **a.** From the 8 solos (monoculture experiments) and 28 duos (pair-wise co-culture experiments), one can calculate the *relative yield R*_*ij*_, quantifying the promotion (positive) or inhibition (negative) impact of species *j* on species *i*. The absolute values shown in the matrix *R* = (*R*_*ij*_) indicate the strengths of promotion and inhibition effects. The sign-pattern of this matrix serves as the ground truth of that of the Jacobian matrix associated with the unknown population dynamics of this microbial community. **b.** Without considering the 8 solos and 28 duos, we analyze the rest steady-state samples. We use the brute-force method to infer the ecological interaction types, i.e., the sign-pattern of the Jacobian matrix. Blue (or red) means inhibition (or promotion) effect of species *j* on species *i*, respectively. 10 signs (labelled by ‘×’) were falsely inferred, 4 signs (grey) are undetermined by the analyzed steady-state samples. **c-d.** The robustness of the inference results in the presence of artificially added noise: 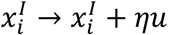, where the random number *u* follows a uniform distribution 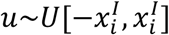, and *η* is the noise level. **c**. At each noise level, we run 50 different realizations. We can see many of inferred *J*_*ij*_ remain their signs in the presence of noise up to noise level *η*=0.3. **d.** At *η* = 0.04, we plot the percentage of unchanged signs for inferred Jacobian matrix in 50 different realizations. The ‘×’ labels correspond to the 10 falsely inferred signs shown in (b). We find that 5 of the 10 falsely inferred results change their signs frequently even when the perturbation is very small, implying that the falsely inferred signs in (b) could be due to measurement noise in the experiments.

Before we apply our inference method, to be fair we remove all those steady states involving one- or two species, and analyse only the remaining 65 steady states. (Note that for *N* = 8, the number of total possible steady states is Ω_max_ = 255. Hence we only use roughly one quarter of the total possible steady states.) During the inference, we first check if the population dynamics of this microbial community can be well described by the GLV model. We find that all the fitted hyperplanes show small *R*^2^, indicating that the GLV model is not appropriate to describe the dynamics of this microbial community (Supplementary Fig. S10b). Hence, we have to aim for inferring the ecological interaction types, without assuming any specific population dynamics model.

Since this microbial community has only eight species, we can use the brute-force algorithm to infer the sign-pattern of the 8×8 Jacobian matrix, i.e., the ecological interaction types between the 8 species (The results of using a heuristic algorithm are similar and described in the Supplementary Fig. 10c). Compared with the ground truth obtained from the relative yield (Fig. 5a), we find that 50 (78.13%) of the 64 signs were correctly inferred, 10 (15.62%) signs were falsely inferred (denoted as ‘×’), and 4 (6.25%) signs cannot be determined (denoted as ‘?’) with the information provided by the 65 steady states (Fig. 5b).

We notice that the *relative yield* of many falsely inferred interactions is weak (with the exception of *R*_Ea,Pch_ and *R*_Ea,Pf_). We conjecture that these errors are caused by noise or measurement errors in the experiments. To test this conjecture, we analyzed the robustness of each inferred *s*_*ij*_ by calculating the percentage of unchanged *s*_*ij*_ after adding perturbations to the samples (Fig. 5c). Similar to adding noise to simulated steady-state data, here we add noise to each non-zero entry 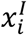 of a sample ***x**^I^* such that 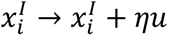 where 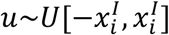. The more robust the inferred results are, the higher the percentage of unchanged signs as *η* is increased. We found that most of the inferred signs were robust: the percentage of unchanged signs remained nearly 80% up to noise level *η* = 0.3 (Fig. 5c). Specifically, Fig. 5d plots the percentage of unchanged signs of the inferred Jacobian matrix when *η* = 0.04. We found that even if the perturbation is very small, 5 of the 10 falsely inferred *s*_*ij*_ in Fig. 5b changed their signs very frequently (blue entries with label ‘×’ in Fig. 5d). In other words, those interactions were very sensitive to noise, suggesting that some falsely inferred signs in Fig. 5b were largely caused by the noise.

#### A synthetic bacterial community of maize roots^44^

There are 7 bacterial species (Ecl, Sma, Cpu, Opi, Ppu, Hfr and Cin) in this community. This dataset consists of in total 8 steady-state samples: 7 sextets and 1 septet. We verified that this community cannot be described by the GLV dynamics (Supplementary Fig. 11).

Using only the 7 sextets (i.e., 7 steady-state samples involving 6 of the 7 species), we inferred the sign-pattern of the Jacobian matrix (Fig. 6a). Based on the sign of *J*_*ij*_, we can predict how the abundance of species-*i* in a microbial community will change, when we add species-*j* to the community. For example, if we add Ecl to a community consisting of the other 6 species (i.e., Sma, Cpu, Opi, Ppu, Hfr and Cin), we predict that the abundance of Sma, Opi, Ppu, Hfr and Cin will increase, while the abundance of Cpu will decrease (first column of Fig. 6b). Note that our prediction only considers the direct ecological interactions between species and ignores the indirect impact among species. Indeed, Ecl promotes Opi, but Ecl also promotes Hfr that inhibits Opi. Hence the net effect of Ecl on Opi is hard to tell without knowing the interaction strengths. Nevertheless, we found that our prediction is consistent with experimental observation (Fig. 6b, first column).

**Figure 6.**
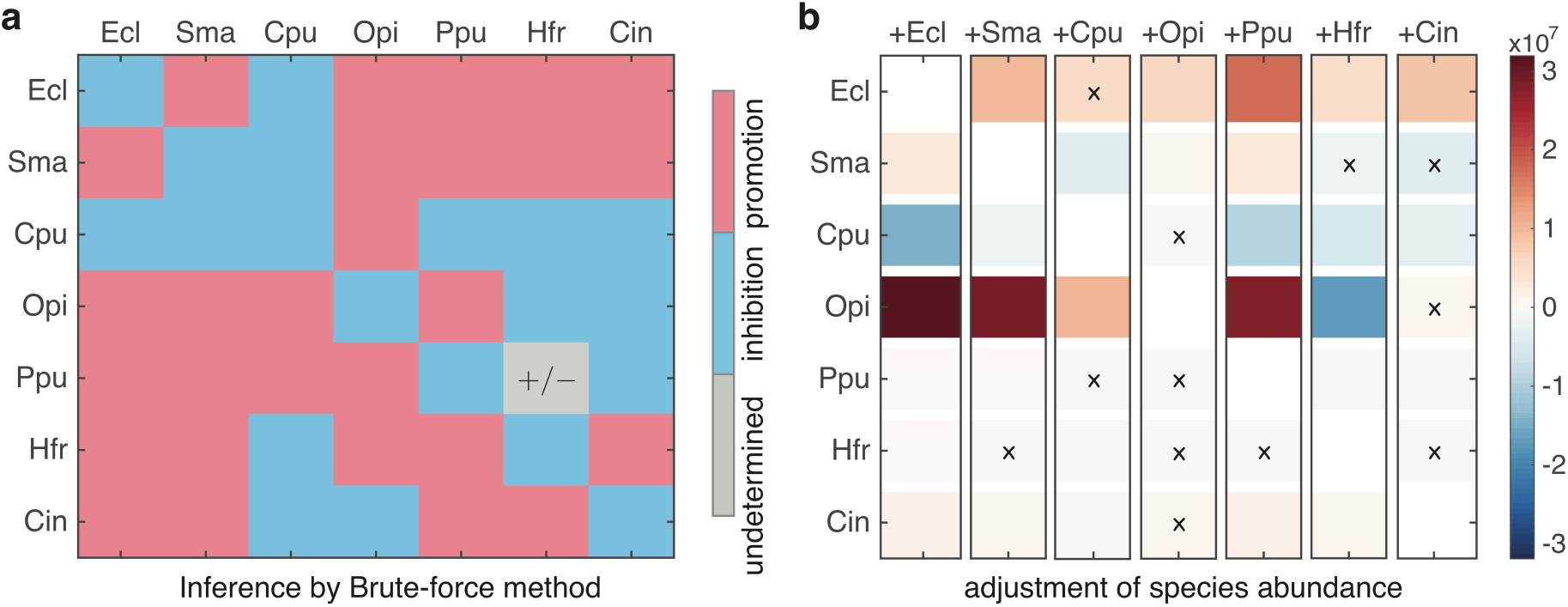
Inferring interaction types in a synthetic community of maize roots with 7 bacterial species. The dataset consists of 7 sextets and 1 septet. **a.** Without considering the 1 septet, we analyze the 7 sextets (steady-state samples involving 6 of the 7 species). We use the brute-force method to infer the ecological interaction types, i.e., the sign-pattern of the Jacobian matrix. Blue (or red) means inhibition (or promotion) effect of species *j* on species *i,* respectively. One sign (grey) are undetermined from the 7 sextets. **b.** The changes of species abundance before and after respectively adding one species into a sextet. Each column corresponds to a sextet (a 6-baterial community), the name of the newly introduced species is marked in the top of each column. Blue (or red) corresponds to the decrement (or increment) of one species after introducing a new species into sextets, respectively. ‘×’ indicates the false prediction. There are in total 12 false predictions.

We then systematically compared our predictions of species abundance changes with experimental observations. There are in total 7 sextets, corresponding to the 7 columns in Fig. 6b. We add the corresponding missing species back to the community, and check the abundance changes of the existing 6 species. There are in total 6×7 = 42 abundance changes. We found that our inferred sign-pattern of the Jacobian matrix (Fig. 6a) can correctly predict 30 of the 42 abundance changes (accuracy ~71.43%). Moreover, for those false predictions, the detailed values of the abundance changes are actually relatively small (comparing to those of correct predictions). Note that we only used 7 steady-samples to infer the interaction types. If we have more steady-state samples available, we assume the prediction accuracy of our method can be further improved.

In the Supplementary Notes 6.3, 6.4, we also demonstrated the application of our method to two additional datasets.

## 3. Discussion

In this work, we developed a new inference method to map the ecological networks of microbial communities using steady-state data. Our method can qualitatively infer ecological interaction types (signs) without specifying any population dynamics model. Furthermore, we show that steady-state data can be used to test if the dynamics of a microbial community can be well described by the classic GLV model. When GLV is found to be adequate, our method can quantitatively infer inter-taxa interaction strengths and the intrinsic growth rates.

The proposed method bears some resemblance to previous network reconstruction methods based on steady-state data^45^. But we emphasize that, unlike the previous methods, our method does not require any perturbations applied to the system nor sufficiently close steady states. For certain microbial communities such as the human gut microbiota, applying perturbations may raise severe ethical and logistical concerns.

Note that our method requires the measurement of steady-state samples and absolute taxon abundances. For systems that are in frequent flux, where steady-state samples are hard to collect, our method is not applicable. Moreover, it fails on analyzing the relative abundance data (see Supplementary Note 2.4 for details). Note that the compositionality of relative abundance profiles also represents a major challenge for inference methods based on temporal data^15,19^. Fortunately, for certain small laboratory-based microbial communities, we can measure the absolute taxon abundances in a variety of ways, e.g., selective plating^46^, quantitative polymerase chain reaction (qPCR)^15,16,47,48^, flow cytometry^49^, and fluorescence in situ hybridization (FISH)^50^. For example, in the study of a synthetic soil microbial community of eight bacterial species^43^, the total cell density was assessed by measuring the optical density and species fractions (relative abundance) were determined by plating on nutrient agar plates. In recent experiments evaluating the dynamics of *Clostridium difficile* infection in mice models^15,16^, two sources of information were combined to measure absolute abundances: (1) data measuring relative abundances of microbes, typically consisting of counts (e.g., high-throughput 16S rRNA sequencing data); and (2) data measuring overall microbial biomass in the sample (e.g., universal 16S rRNA qPCR).

In contrast to the difficulties encountered in attempts to enhance the informativeness of temporal data that are often used to infer ecological networks of microbial communities, the informativeness of independent steady-state data can be enhanced by simply collecting more steady-state samples with distinct taxa collection (For host-associated microbial communities, this can be achieved by collecting steady-state samples from different hosts). Our numerical analysis suggests that the minimal number of samples with distinct taxa collections required for robust inference scales linearly with the taxon richness of the microbial community. Our analysis of experimental data from a small synthetic microbial community of eight species shows that collecting roughly one quarter of the total possible samples is enough to obtain a reasonably accurate inference. Furthermore, our numerical results suggest that this proportion can be significantly lower for larger microbial communities.

This blessing of dimensionality suggests that our method holds great promise for inferring the ecological networks of large and complex microbial communities, such as the human gut microbiota. There are two more encouraging facts that support this idea. First of all, it has been shown that the composition of the human gut microbiome remains stable for months and possibly even years until a major perturbation occurs through either antibiotic administration or drastic dietary changes^51-54^. The striking stability and resilience of human gut microbiota suggest that the collected samples very likely represent the steady states of the gut microbial ecosystem. Second, for healthy adults the gut microbiota displays remarkable universal ecological dynamics^55^ across different individuals. This universality of ecological dynamics suggests that microbial abundance profiles of steady-state samples collected from different healthy individuals can be roughly considered as steady states of a conserved “universal gut dynamical” ecosystem and hence can be used to infer its underlying ecological network. Despite the encouraging facts, we emphasize that there are still many challenges in applying our method to infer the ecological network of the human gut microbiota. For example, the assumption of invariant ecological interaction types (i.e., promotion, inhibition, or neutral) between any two taxa needs to be carefully verified. Moreover, our method requires the measurement of absolute abundances of taxa.

We expect that additional insights into microbial ecosystems will emerge from a comprehensive understanding of their ecological networks. Indeed, inferring ecological networks using the method developed here will enable enhanced investigation of the stability^56^ and assembly rules^57^ of microbial communities as well as facilitate the design of personalized microbe-based cocktails to treat diseases related to microbial dysbiosis^9,10^.

## Acknowledgements

This work is supported in part by the John Templeton Foundation (Award number 51977). We thank Drs. Gabe Billings and Brigid Davis for insightful comments on the manuscript. We thank Drs. Joseph Nathaniel Paulson, Michael T. Mee, Harris H. Wang, Francesco Carrara and Carsten F. Dormann for kindly providing their experimental datasets. We thank Dr. Liang Tian for discussions.

## Contributions

Y.-Y.L conceived the project. Y.-Y.L and M.T.A. designed the project. Y.X. and M.T.A. did the analytical calculations. Y.X. did the numerical simulations and analyzed the empirical data. All authors analyzed the results. Y.-Y.L., Y.X. and M.T.A. wrote the manuscript. All authors edited the manuscript.

## Author Information

The authors declare no competing financial interests.

## 1. Theoretical Basis For Inferring Ecological Interactions

Here we formulate and prove two theorems (Theorems 1 and 2) that characterize the conditions for inferring the presence/absence or type (positive, negative or neutral) of ecological interactions in a microbial community using steady-state data. These theorems provide the basis for the inference methods described in Supplementary Note 2.

### 1.1. Notation

We use bold letters like *x* to denote vectors, and capitals like *J* to denote matrices. The *i*-th element of the vector *x* is denoted by *x*_*i*_. Similarly, *J*_*i*_ denotes the *i*-th row of matrix *J*, and *J*_*ij*_ denotes the (*i, j*)-th element of matrix *J*. For a matrix 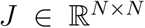, we denote by *S* = (***s***_*ij*_) = sign(*J*) ∈ {−1, 0,1}^*N×N*^ its sign-pattern, where ***s***_*ij*_ = sign(*J*_*ij*_). Similarly, we denote by *Z* =(*z*_*ij*_) ∈ {0,1}^*N×N*^ its zero-pattern, where *Z*_*ij*_ = |*s*_*ij*_|.

### 1.2. Preliminaries

Consider a microbial community of *N* different taxa. Let *x*_*i*_*(t)* denote the absolute abundance of taxon *i* at time *t*. Suppose the temporal evolution of the taxa abundances are described by a generic population dynamics model taking the form of a set of ordinary differential equations (ODEs):

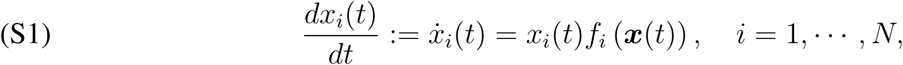

where 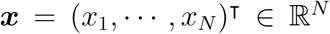 is the state vector and 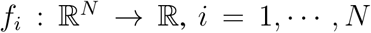, are some non-zero *meromorphic* functions—that is, the quotient of two analytical functions of *x*.

#### Remark 1.

Typical examples of meromorphic functions are in the form of the quotient of two polynomials. By specifying these meromorphic functions, system (S1) can take the form of many classical population dynamics models [1, 2, 3, 4, 5, 6]. The assumption that all *f*_*i*_*(x)*’s are some non-zero meromorphic functions has a useful consequence, since meromorphic functions have the *generic* properties inherited from analytic functions [7]. This implies, for example, that since the *f*_*i*_’s are not identically zero, they can be zero only on a zero-measure set of their domain 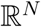.

A steady-state dataset 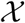 is a collection of *N*-dimensional vectors 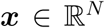 corresponding to the measured equilibria of Eq. (S1). Each element of 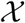 is called a steady-state sample, or just a sample. We will denote a sample as 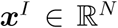, where its taxon *index set* 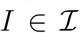 determines which taxa are present. Here 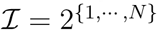 is the set of all possible subsets of {1, ·, *N*}. For example, 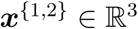 is a sample of a community with three taxa, and in this sample only taxon 1 and taxon 2 are present.

Consider now the subset 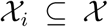 of all samples containing taxon *i*, so that *f*_*i*_*(x)* = 0 for all 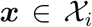. Then, for any two samples 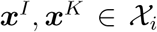, applying the the mean value theorem for multivariable functions, we obtain

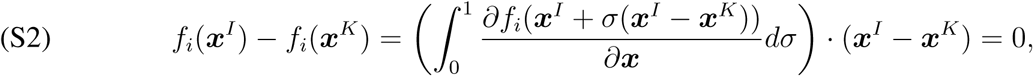

where ‘·’ denotes the inner product between vectors in 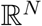. Let 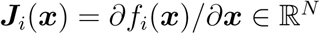 be the *i*-th row of the Jacobian matrix *J(x)* = (*J_ij_(x))* = (*∂f*_*i*_*(x)/∂*_*x*_*j*__) and let us introduce the notation

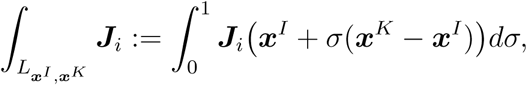

where *L*_*x*^*I*^_,_*x*^*K*^_ denotes the line segment joining the points *x^I^* and *x^K^* in 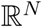. With this notation, Eq. (S2) can be rewritten more compactly as

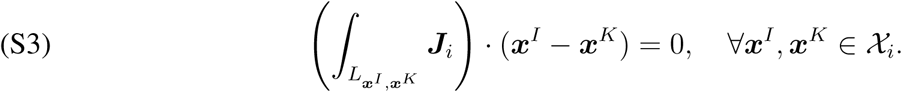

The above equation implies that the difference of any two samples {***x**^I^, x^K^*} sharing taxon *i* constrains the integral of ***J***_*i*_ over the line segment joining them ***x**^I^* − ***x**^K^*.

In this work, we consider that the ecological interactions in a microbial community are encoded in the Jacobian matrix 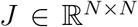 of its population dynamics. More precisely, we assume that the j-th taxon directly impacts the *i*-th one iff the function *J*_*ij*_(*x*) ≢ 0. Notice that this condition is well defined because *J*_*ij*_(*x*) is a meromorphic function. Further, an ecological interaction is inhibitory iff *J*_*ij*_(*x*) < 0 and excitatory iff *J*_*ij*_(*x*) > 0.

Inferring the absence or presence of interactions is equivalent to inferring the zero-pattern of the Jacobian matrix, recovering the topology of the ecological network underlying the microbial community. Furthermore, inferring the type of interactions (inhibitory, excitatory or null) is equivalent to inferring the sign-pattern of the Jacobian matrix. Analyzing the implications of Eq. (S3) will be the basis for inferring these two properties of the Jacobian matrix from the steady-state samples 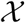, as we next show.

### 1.3. Inferring the zero-pattern

Let 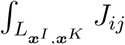 denote the *j*-th entry of the vector 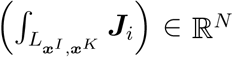. To infer the zero-pattern of the Jacobian matrix, we make the following assumption:

#### Assumption 1.

The condition 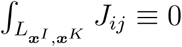 holds if and only if *J*_*ij*_ ≡ 0 for all *i,j* = 1,…, *N*.

#### Remark 2.

a. Assumption 1 is a *necessary condition* to recover the zero-pattern of ***J***_*i*_, *i* = 1,…, *N*, from steady-state samples 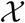 regardless of the algorithm used for the inference. Indeed, if 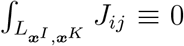 but *J*_*ij*_ ≢ 0, then it is impossible to distinguish from the samples 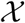 if either *J_ij_(x)* ≡ 0 or *J_ij_(x)* ≢ 0, as both conditions would satisfy Eq. (S3).
b. Assumption 1 is generically satisfied for most functions *f*_*i*_*(x)* used in population dynamics models. More precisely, notice how the condition 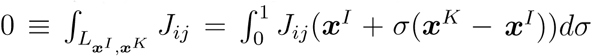 requires that the positive and negative areas below *J*_*ij*_(***x**^I^* + σ(***x**^K^* − ***x**^I^*)) cancel exactly when plotted as a function of *σ.* Such condition is not *generic.* This means that if the above equation holds for some particular *f*_*i*_ and its corresponding *J*_*ij*_, then there exists an infinitesimal deformation 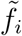 of *f*_*i*_ such that the areas of the corresponding 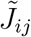 as a function of *σ* do not cancel out anymore.

For each sample pair *{x^I^, x^K^*}, let’s denote by 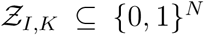 the set of zero-patterns of all vectors orthogonal to ***x**^I^ − x^K^*. Then we obtain the following result:

#### Theorem 1.

Let *Z*_*i*_ ∈ {0,1}^*N*^ be the zero-pattern of ***J***_*i*_. Then, under Assumption 1, we have that

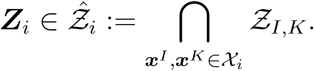

*Proof.* From Assumption 1, we conclude that *Z*_*i*_ and the zero-pattern of 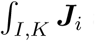 are identical for all 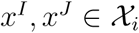. Then Eq. (S3) implies that the 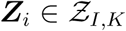 for all 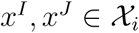. This directly implies that 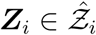. □

#### Remark 3.

a. Note that 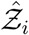 will always contain at least two elements: a trivial one 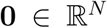 and a nontrivial one 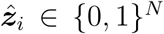. Therefore, to distinguish which of these two zero-patterns is the true zero-pattern of *J*_*i*_, it is necessary to a-priori know the existence of at least one nonzero interaction.
b. Theorem 1 together with Remark 3.a provide the basis of a computational method to infer the zero-pattern of the Jacobian matrix, since the sets 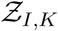 can be computed from the steady-state data 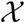.

#### Example 1.

Consider a toy model with two taxa:

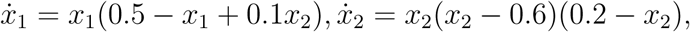

where the Jacobian matrix is

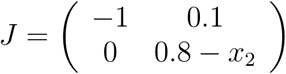

Note that the sign of *J*_22_ depends on the value of *x*_2_. Supplementary Fig. 1b shows that *J*_22_ indeed changes its sign from positive to negative during the growth process (Supplementary Fig. 1a).

In the absence of measurement noise, we can successfully infer the zero-pattern of *J* (Supplementary Fig. 1c,d). Supplementary Fig. 1c shows that, according to the position of ***x***^{1,2}^ − ***x***^{1}^, the green line that is orthogonal to the red line cannot yield a zero entry for ***J***_1_, implying that *J*_11_ ≠ 0 and *J*_12_ ≠ 0. This is consistent with the ground truth. Supplementary Fig. 1d shows that for ***J***_2_, the green line that is orthogonal to the blue line exactly locates on the axis of *x*_2_, implying that *J*_21_ = 0 and *J*_22_ ≠ 0, which is also consistent with the ground truth.

In the presence of measurement noise, the angle between the *x*_1_-axis (or the *x*_2_-axis) and the green line can be used to determine if *J*_*ij*_ = 0 or not (Supplementary Fig. 1e,f). For example, when the noise level (*η*) is 0.1, in Supplementary Fig. 1e the angle between the *x*_1_-axis and the green line is large enough and we can safely conclude that *J*_12_ ≠ 0. By contrast, the green line deviates only slightly from the *x*_2_-axis and these deviations are randomly distributed and have zero mean (Supplementary Fig. 1f). Therefore, we can choose a threshold value *θ* such that if the absolute value of the average deviation angle over different measurements is smaller than *θ*, we conclude that *J*_21_ = 0. Notice that this method will infer very weak interactions as zero, but it still offers a pragmatic approach to infer strong interactions.

### 1.4. Inferring the sign-pattern

In order to infer the sign-pattern, we assume:

#### Assumption 2.

The nature of ecological interactions (i.e., parasitism, commensalism, mutualism, amensalism or competition) between any two taxa does not vary over the collected steady-state samples.

Note that Assumption 2 is actually necessary to infer the ecological interaction types. If those interaction types vary from samples to samples, then the inference becomes an ill-defined problem because we have a “moving target” and different subsets of steady-state samples will offer different answers on the interaction types.

#### Remark 4.

Assumption 2 has the following consequences:

a. For *x* in the positive orthant of 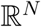, each element *J*_*ij*_(*x*) of the vector ***J***_*i*_(*x*) is either uniformly negative, uniformly zero or uniformly positive. Indeed, if and only if this condition is satisfied, the sign-pattern of the Jacobian, given by 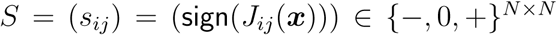, is constant.
b. The sign-pattern of the vectors 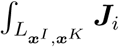 is the same for all 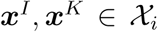, and it coincides with the sign-pattern of ***J***_*i*_.

With Assumption 2, next we show that the vector (***x**^I^* − ***x**^K^*) constrains enough the possible sign-pattern of the vector ***J***_*i*_ so that we can infer it. Let us associate each orthant of 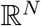 with its corresponding sign-pattern, that is, a vector in {−, 0, +}^*N*^. There are exactly 3^*N*^ vectors in {−, 0, +}^*N*^.

#### Example 2.

{−, 0, +}^2^ has 9 sign-patterns:

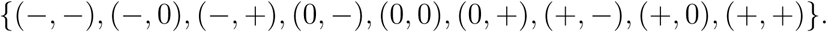

And {− 1, 0,1}^3^ has 27 sign-patterns.

In order to show how the vector (***x***^*I*^ − ***x***^*K*^) constrains the sign-pattern of ***J***_*i*_, we start by discussing the following elementary example:

#### Example 3.

Consider *N* = 2 and three steady-state samples *x*^{1}^, *x*^{2}^ and *x*^{1,2}^. From Eq. (S3), we obtain:

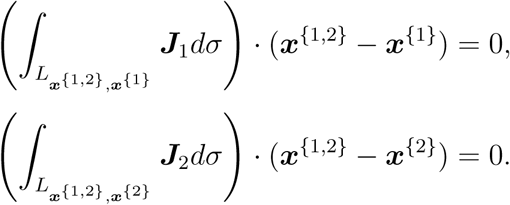

These equations imply that 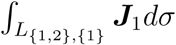 is orthogonal to the line *L*_{1,2},{1}_, and that 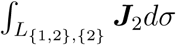 is orthogonal to the line *L*_{1,2},{2}_. Further, as discussed in Remark 4.b, recall that Assumption 2 implies `sign`(∫ *J*_*j*_)= `sign`(*J*_*j*_). Hence, `sign`(***J***_i_) is one of the sign-patterns corresponding to the line orthogonal to *L*_{*i*,2},{*i*}_, see Supplementary Fig. 2.

As a concrete example showing how the above equations can be used to infer the sign-pattern of the Jacobian matrix, consider the following ecological dynamics with the so-called Holling Type II functional response [2]:

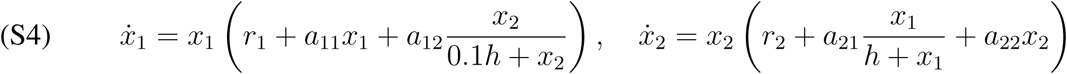

Here 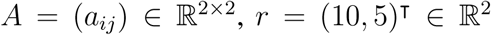 and *h* = 1 are parameters. Importantly, notice that `sign`(*J*) = `sign`(*A*).

Next we focus on inferring `sign`(*J*_1_), as the same procedure can be applied to infer `sign`(*J*_2_). For illustration, we choose *a*_21_ = −1, *a*_22_ = −1 and then consider two cases for the remaining two parameters (*a*_11_,*a*_12_):

Case 1. For *a*_11_ = −1 and *a*_12_ = 1, the feasible steady states^1^ of Eq. (S4) are

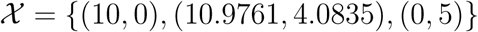

see Supplementary Fig. 2a. In order to infer `sign`(***J***_1_), we focus on the line *L*_{1},{1,2}_ connecting the samples ***x***^{1}^ = (10,0) and ***x***^{1,2}^ = (10.9761,4.0835) where taxon 1 is present. The line orthogonal to *L*_{1},{1,2}_ (shown in green) determines the possible sign-patterns for ***J***_1_. The sign-patterns corresponding to this orthogonal line are 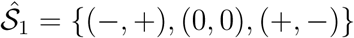 and notice that 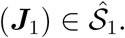.

Case 2. For *a*_11_ = −1 and *a*_12_ = −1, the feasible steady states of Eq. (S4) are

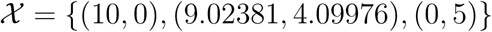

see Supplementary Fig. 2b. We focus on the line orthogonal to *L*_{1},{1,2}_ (shown in green) and obtain its corresponding sign-patterns 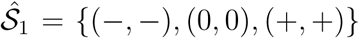. Again, notice that 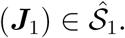.

In the general case of *N* taxa we have the following result:

#### Theorem 2.

Let ***S***_*I,K*_ ⊆ {−, 0, +}^*N*^ be the set of all sign-patterns associated with the vectors orthogonal to (*x*^*I*^ − ***x***^*K*^) and define 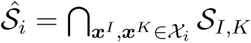. Then `sign`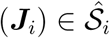.

*Proof.* From Assumption 2, we know that `sign`(***J***_i_) = `sign`(∫_*I,K*_ *J*_*i*_) for all ***x**^I^, x^K^* ∈ *χ*_*i*_. Then Eq. (S3) implies that 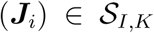 for all pairs ***x***^*I*^,*x*^*K*^ ∈ *χ*_*i*_. Thus, `sign`(*J*_*i*_) must belong to the intersection of all 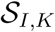, implying that `sign`(*J*_*i*_) belongs to the sign-patterns shared by all 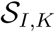.

#### Remark 5.

a. In Theorem 2, in order to check if there is a vector with sign-pattern 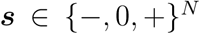 orthogonal to a given vector (***x***^*I*^ − ***x***^*K*^), we can check if the following linear program has a solution:

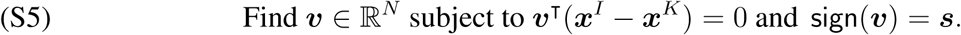 Note that the condition `sign`(*v*) = *s* can be encoded as a set of equalities/inequalities of the form *(v*_*i*_ = 0, *v*_*i*_ < 0, *v*_*i*_ > 0} corresponding to the the cases {***s***_*i*_ = 0, ***s***_*i*_ = −1, ***s***_*i*_ = 1}. in Supplementary Note 2.3.1, we will show that it is possible to solve this problem more efficiently using the notion of sign-satisfaction.
b. From a geometrical viewpoint, the vectors 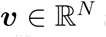 satisfying *v*^T^(***x***^*I*^ − ***x***^*K*^) = 0 correspond to the hyperplane with normal vector (***x***^*I*^ − ***x***^*K*^). Thus, the set of sign-patterns of those vectors *v* (i.e., the sign-patterns in the set 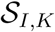) corresponds to the orthants of 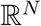 crossed by this hyperplane. Consequently, 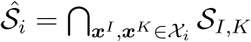 corresponds to those orthants of 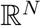 crossed by *all* hyperplanes orthogonal to (***x***^*I*^ − ***x***^*K*^) for all 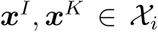.
c. Note that 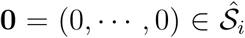 always. Additionally, there is always at least three admissible sign-patterns in 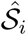, that is 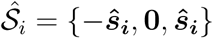 for some 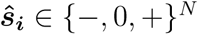.
d. A consequence of Remark 5.b above is that the steady-state data alone cannot be informative enough to determine a unique sign-pattern for ***J***_*i*_. The number *m* of sign-patterns (candidate solutions) in 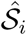 depends on the number and informativeness of the samples 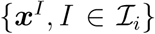 in the steady-state dataset *χ*. Indeed, if *m* sign-patterns are in 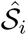, only by providing the sign of exactly ∟*m*/3⌋ non-zero entries of ***J***_*i*_ as prior information, we can univocally infer `sign`(***J***_*i*_). For instance, in the case of *N* =2 taxa in Example 2 where *m* = 3, we need to provide the sign of only one non-zero entry of ***J***_*i*_, and that will let us infer the sign of the other entry.

To conclude this subsection, we discuss some implications of Assumption 2 on the steady-state samples *χ* that can be observed in the microbial community. Let 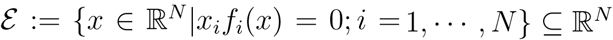 denote the set of equilibria of Eq. (S1).

#### Definition 1.

System (S1) is said to have *true multi-stability* if *ɛ* contains at least two isolated sets (i.e., two sets that don’t intersect) of interior equilibria.

A particular case of true multi-stability is when the system or any subsystem (composed of a particular subset of taxa) exhibits multiple interior equilibria, where all the involved taxa have positive abundances.

#### Proposition 1.

If Assumption 2 is satisfied then there is no true multi-stability.

***Proof.*** We argue by contradiction. Suppose that Assumption 2 is satisfied but the system has true multi-stability. Let ***x***^*I*^ and ***x***^*K*^ be two interior equilibria belonging to two different isolated sets in *ɛ*. Suppose they are interior with respect to the *i*-th taxon. Now consider the scalar function *p(σ)* = *f*_*i*_(***x***^*I*^ + *σ*(*x*^*K*^ − *x*^*I*^)). Note that *p(σ)* is a non-zero meromorphic function because ***x***^*I*^ and ***x***^*K*^ belong to different isolated sets of equilibria. Furthermore, *p*(0) = *p*(1) = 0 as both ***x***^*I*^ and ***x***^*K*^ are interior equilibria for *f*_*i*_*(x)*. Together, this implies that the slope of *p(σ)* needs to change sign at least once. Since the slope of *p(σ)* equals the Jacobian of *f*_*i*_ in the direction of the vector *x*^*I*^ − *x*^*K*^, this implies that the Jacobian changes sign, contradicting Assumption 2.

#### Remark 6.

a. Proposition 1 actually provides a simple criterion to falsify Assumption 2. Namely, if a microbial community displays true multi-stability, then Assumption 2 is invalid, i.e., the sign-pattern of its Jacobian matrix is not constant. In practice, we can detect the presence of true multi-stability in the available samples 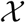 (e.g., two or more steady-state samples have the same collection of present taxa but totally different abundance profiles). If multi-stability is detected, then we know immediately that Assumption 2 is invalid. If multi-stability is not detected, then at least Assumption 2 is consistent with the collected steady-state samples. In short, by introducing a criterion to falsify our assumption, we significantly enhance the applicability of our method for inferring interaction types.
b. In case the sign-pattern of the Jacobian matrix is not constant but the steady-state samples were still collected from a microbial community under the same or similar environmental conditions (e.g., nutrient availability), we can interpret our inferred sign-patterns as the overall inhibition or promotion effect between different taxa during the transitions between steady states. To see this point, note that regardless of the sign-pattern of the Jacobian matrix is constant or not, our method can correctly infer the sign of 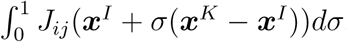, which reflects an overall impact (inhibition or promotion) of taxon *j* on taxon *i* during the transition from the steady state *x*^*I*^ (*σ* = 0) to *x*^*K*^ (*σ* = 1).

#### Example 4.

To illustrate Remark 6.b, let’s consider a toy model of twu species *X* and *Y*. Each species has a per capita growth rate that is modulated by its mutualistic partner as well as the resource. The population dynamics of this toy model is given by

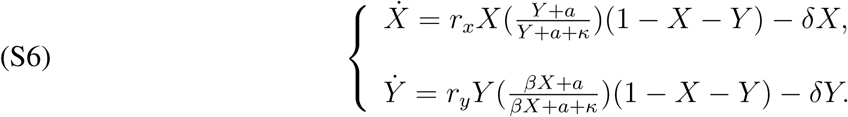

Here *r*_*x*_ and *r*_*y*_ are the growth rates of the species, *a* is the amount of resource, *δ* is the death rate, *κ* is an effective Monod constant, and *β* > 0 quantifies the asymmetry of benefit that each species receives from its partner. The elements of the Jacobian matrix of this community are given by

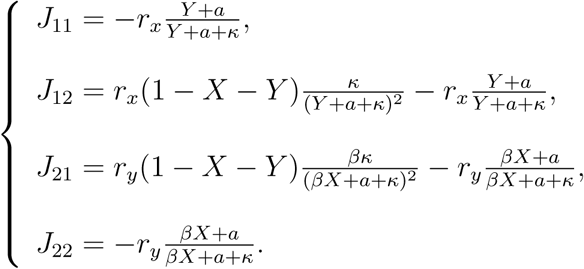

Note that *J*_11_ and *J*_22_ are always negative, while *J*_12_ and *J*_21_ may change their signs depending on the particular abundances of *X* and *Y*, as well as the model parameters. This model captures the transition between the different regimes of ecological interaction depending on the amount of resource (determined by *a*). Indeed, Supplementary Fig. 3a shows that there are three regimes with different inter-species interactions starting from mutualism and then leading to parasitism and competition. Here, the overall or “effective” interaction types are determined by comparing the difference of steady states between monocultures (dashed lines in Supplementary Fig. 3a) and co-cultures (solid lines in Supplementary Fig. 3a). This allow us to calculate the relative yield, which indicates the promotion or inhibition impact between two taxa. For this model, because that Jacobian may change its sign-pattern over time, the sign of relative yields can be interpreted as the effective impact between two taxa, denoted as *J*_eff_ (Supplementary Fig. 3a).

We now apply our method using steady states of this model. Supplementary Fig. 3b-d shows the diagrams of our inference method under different resource amounts. We found that the inferred inter-species interaction types are consistent with the ground truth (shown in Supplementary Fig. 3a). Supplementary Fig. 3e-g shows the value of *J*_12_(***x***^{1,2}^ + σ(***x***^{2}^ − ***x***^{1,2}^))*dσ* and *J*_21_(***x***^{1,2}^ + *σ*(***x***^{2}^ − ***x***^{1,2}^))*dσ* as a function of *σ* corresponding to the transition between ***x***^{1,2}^ *(σ* = 0) and ***x***^{2}^ *(σ* =1). The shade areas in this figure denote the value of 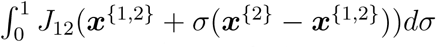 and 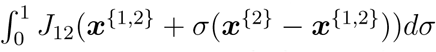. For example, when *a* = 0.15, both 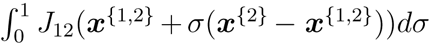 and 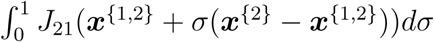 are positive (shaded areas in Supplementary Fig. 3e). Although *J*_12_ and *J*_21_ display both negative and positive values as *σ* changes, the positive *J*_12_ and *J*_21_ dominate in the transition between two steady states. Hence, overall taxon *X* and taxon *Y* are mutualistic. Supplementary Fig. 3f,g shows the result of this analysis applied to *J*_12_ and *J*_21_ for *a* = 0.2 and *a* = 0.5.

## 2. BRUTE-FORCE AND HEORISTIC ALGORITHMS

Here we introduce the methodology for inferring the zero- or sign-patterns of the Jacobian matrix associated with the population dynamics of a microbial community. In essence, the inference of the zero-pattern is similar to the inference of the sign-pattern. Indeed, the only difference is that the former doesn’t care if the non-zero values are positive or negative. This implies that the complexity for inferring the network topology and interaction types are roughly the same. Here, for simplicity we describe the algorithms for inferring sign-patterns. All the algorithms (and pseudo codes) can be easily modified to infer the zero-pattern.

### 2.1. Brute-force algorithm

Theorems 1 and 2, together with Remarks 3 and 5 can be used to construct an algorithm to obtain all admissible sign-patterns for given steady-state data. Indeed, by enumerating all possible sign-patterns, we can use the liner program in Eq. (S5) to check if each of the possible 3^*N*^ sign-patterns is admissible for taxon i, see Algorithm 1.

#### Algorithm 1 A brute-force algorithm to compute 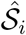

~~~
**Input:**
  The collection of matrices *M*_*i*_, being the difference between all two samples containing species 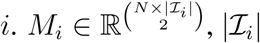 is the nomber of samples
**Output:**
  The sign-pattern set of 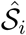
1: 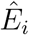 ← Enomeration of all the possible combinations {−, 0, +}^*N*^
2: **for** each *j*-th row in *M*_*i*_ **do**
3:   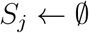
4:   **for** each *k*th-subset in 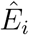 **do**
5:    if find 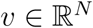 subject to *v*^⊤^*M*_*i*_[*j*,:] = 0 and 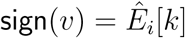 **then**
6:     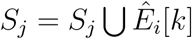
7:    **end if**
8:   **end for**
9:   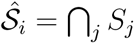
10: **end for**
~~~

Below, we illustrate the application of the brute-force algorithm for a microbial community with *N* = 3 taxa.

#### Example 5.

Here we consider the case of a microbial community with *N* = 3 taxa and population dynamics given by the so-called Crowley-Martin functional response [5]. The ODEs are:

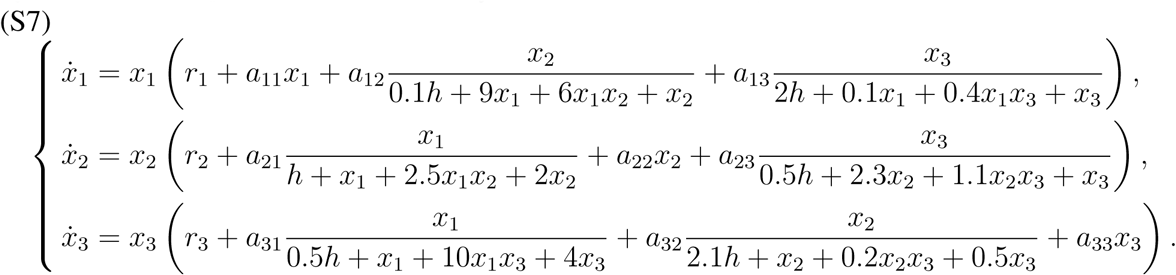

We set the parameters *r*_1_ = 1, *r*_2_ = 5, *r*_3_ = 1.5, *h* = 0.2 and

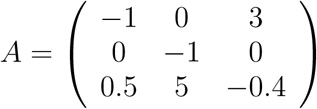

Notice again that `sign`(*J*) = `sign`(*A*). We focus on reconstructing `sign`(***J***_1_), as the same procedure applies to the other taxa. With the given parameters, the feasible steady states of Eq. (S7) where taxon 1 is present are:

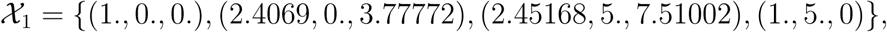

constituting the available steady-state samples for taxon 1. We apply Algorithm 1 to this dataset obtaining

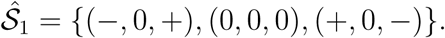

Note that 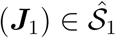. Providing for example `sign`(*J*_11_) < 0 as prior information, we correctly infer that `sign`(*J*_1_) = (−, 0, +).

### 2.2. Computational complexity of the brute-force algorithm

Algorithm 1 strongly relies on the enumeration of all 3^*N*^ possible sign-patterns in 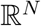, since it needs to test if each one of them is admissible for the given data. If the set 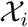 has *n*_*i*_ elements, there will be *n*_*i*_(*n*_*i*_ − 1)/2 vectors of the form ***x***^*I*^ − ***x***^*K*^ with 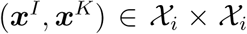. According to Algorithm 1, for each of those vectors and each of the possible 3^*N*^ sign-patterns, we will need to run the linear program (S5) to check if there is an orthogonal vector with the desired sign-pattern.

If we assume that the linear program can be solved with *N* operations, then, for each taxon, Algorithm 1 requires to perform a number of operations in the order of

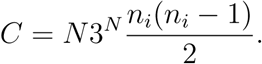

If we have more than two samples (i.e., *n*_*i*_ > 2), then *n*_*i*_(*n*_*i*_ − 1)/2 > 1 and consequently *C* > *N*3^*N*^. Hence, for 100 taxa, we will need to perform at least 5.19 × 10^49^ operations for the reconstruction of each taxon —which is a number with the same order of magnitude as the number of atoms in Earth. Further, the linear programming used in the brute-force method can also be time consuming even for a small microbial community with *N* ~ 10. Consequently, applying the enumeration procedure is only reasonable for a community with *N* ~ 10, since in this case only around 10^6^ operations are needed to infer the sign-pattern of the Jacobian corresponding to each taxon.

The above two limitations motivated us to develop a more efficient reconstruction method. This method has two main ingredients. First, a graph-based approach to quickly check whether a region can be crossed by a hyperplane, circumventing the need to solve the linear program. Second, an heuristic algorithm efficiently explore the solution space and to infer the ecological interaction types.

### 2.3. Inference using the heuristic algorithm

In practice, for large microbial communities with unknown dynamics, the inference of ecological interactions according to Algorithm 1 has two major drawbacks:

1. Checking if an orthant is crossed by a given hyperplane using the linear program of Eq. (S5) is computationally expensive.
2. The number of orthants that is necessary to check (i.e., the solution space) increases as 3^N^, that is, exponentially in the number of taxa.

To circumvent the first drawback, we introduce an alternative method based on the notion of sign-satisfaction. To address the second challenge, we propose an heuristic algorithm with user-defined time complexity to infer the sign-pattern of ***J**_i_.*

#### 2.3.1. Formulating the sign-satisfaction problem

Consider a real-valued vector 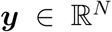. Then, solving the linear program Eq. (S5) is equivalent to solving the following sign-satisfaction problem:

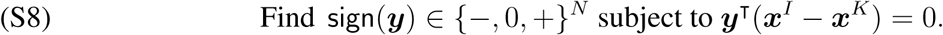

Notice that from a geometrical viewpoint, solving Eq. (S8) is just finding the orthants of 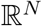 crossed by the hyperplane orthogonal to (***x***^*I*^ − ***x***^*K*^).

In the next example, we illustrate how the sign-satisfaction formulation allows us to quickly discard orthants of 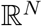 that cannot be crossed by such hyperplane:

##### Example 6.

In vector form, Eq. (S3) in Example 5 can be written as

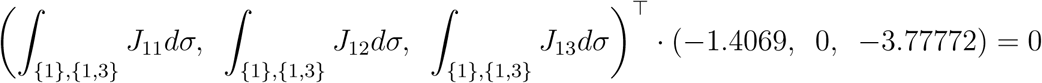

if we take two samples sharing taxon 1. Thus, the sign-satisfaction for Example 5 can be written as

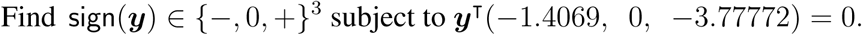

Note that, for example, the choice `sign`(***y***) = (−, 0, −) cannot satisfy the above condition regardless of the particular value of *y*, because the inner product is the sum of two positive numbers, which can never be zero.

A systematic method to extend the above example and solve the sign-satisfaction problem is discussed next.

#### 2.3.2. A graph-based approach to solving the sign-satisfaction problem

We illustrate the basic idea using a small example, and then discuss the general case.

In Example 6, the sign-satisfaction problem required that

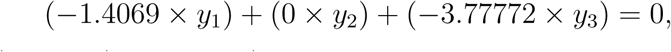

where `sign`(*y*_1_,*y*_2_,*y*_3_) = `sign`(*J*_11_,*J*_12_,*J*_13_). We map the above equation to the sign-satisfaction graph in Supplementary Fig. 4, where each element of ***J***_1_ corresponds to a column and each element of `sign`(***J***_1_) has three possibilities (i.e., ‘−’,‘0’ or ‘+’). Each node in Supplementary Fig. 4 is divided in two parts: the left is an entry of `sign`(***x***^*I*^ − ***x***^*K*^) and the right is an entry of `sign`(***J***_1_). The color of each node encodes the sign of the product of left and right parts: grey is zero, red is positive and blue is negative. Next we introduce edges starting from each node and pointing to all nodes located in the next column to its right. With this formulation, the solutions to the sign-satisfaction problem (S8) reduces to finding the paths in the sign-satisfaction graph that satisfy one of the following two conditions:

i. the path contains red (representing positive values) and blue (representing negative values) nodes simultaneously, or
ii. the path contains only gray (representing zero values) nodes.

The above two conditions guarantee that the sum of the product of `sign`(*J*_*ij*_) and (***x***^*I*^ − ***x***^*K*^) can be zero. At this step, it is also useful to introduce the prior information that is available, such as *J*_*ii*_ < 0, allowing us to collapse columns of nodes in the sign-satisfaction graph to single nodes (Supplementary Fig. 4).

In a general case, for a given (***x***^*I*^ − ***x***^*K*^), the construction of the sign-satisfaction graph is as follows:

1. the graph consists of N columns, with each column having three nodes;
2. each node in the graph is divided into two parts: the left correspond to an entry of `sign`(***x***^*I*^ − ***x***^*K*^) and the right to an entry of `sign`(***J*_*j*_**);
3. each node is colored according to the sign of the product of the left and right parts: zero is grey, positive is red and negative is blue;
4. directed edges are included from a node to all the nodes in next column.

Finally, a solution to the sign-satisfaction problem of Eq. (S8) corresponds to a path from the first column to *N*-th column satisfying either condition (i) or (ii) listed above. In such case, a possible sign-pattern of ***J***_1_ consists of the sign in the right part of each node in the path. For instance, the paths with yellow directed edges in Supplementary Fig. 4 correspond to the possible sign-pattern of ***J**_1_,* i.e., (−,+,+), (−, −, +) and (−, 0, +).

By using the sign-satisfaction graph, it is very efficient to test if the hyperplane orthogonal to (***x**^1^* − ***x***^*K*^) crosses some orthants of 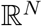, because it reduces to checking if its corresponding vector in {−, 0, +}^*N*^ satisfies either condition (i) or (ii). However, finding all orthants crossed by such orthogonal hyperplane remains challenging, since the sign-satisfaction graph did not decrease the dimension of the solution space (that remains with exponential size 3^*N*^). To address this issue, next we introduce a method to efficiently sample paths in the sign-satisfaction graph.

#### 2.3.3. Use the intersection of hyperplanes to sample paths in the sign-satisfaction graph

As discussed before, with the sign-satisfaction graph the solution space is still exponential (with size 3^*N*−1^, where the term *N* − 1 comes from assuming we know that *J*_*ii*_ < 0 as prior information). One possibility to circumvent this problem would be to randomly sample paths in the sign-satisfaction graph and check if they satisfy conditions (i) or (ii). This would not work, however, since the probability of sampling the true “`sign`(***J***_*i*_)” is only *X*/3^*N*−1^ − where *X* is the number of sampled paths − and this probability approaches zero as *N* increases. To alleviate this problem, next we propose a method to sample paths in the sign-satisfaction graph with certain preference.

This method has four steps and depends on an user-defined parameter Ψ > 1 specifying the number of times the procedure is repeated:

1. **Construct the matrix of the difference of all the sample pairs.** Consider the set of all vectors 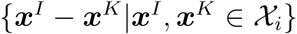. Let 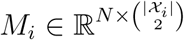 be a matrix constructed by stacking all the 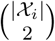 vectors, where 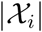 is the number of samples containing taxon *i.* By construction, each column of *M*_*i*_ is the normal vector of a hyperplane orthogonal to the difference of the corresponding sample pair.
2. **Randomly sample (*N* − 1) hyperplanes.** Choose randomly *N* − 1 columns from *M*_*i*_.
3. **Find the intersection of the (*N* − 1) sampled hyperplanes to obtain an intersection line.** This can be done by finding the kernel of the matrix obtained by stacking the chosen columns. Note that the randomly sampled (*N* − 1) hyperplanes not always intersect in a line, because some hyperplanes might be parallel. However, this situation is non-generic in 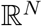. Thus, if the randomly sampled hyperplanes do no intersect as a line, we return to step 2 and choose a new subset of columns.
4. **Count how many hyperplanes cross the region of the intersection line using the sign-satisfaction graph.** The sign-pattern of this intersection line represent the three orthants in 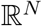 crossed by all those (*N* − 1) hyperplanes. For the remaining hyperplanes in *M*_*i*_ (i.e., the rest of the columns in *M*_*i*_), let 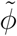 be the number of those hyperplanes that cross these three orthants. We normalize 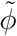 using 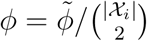, so that *ϕ* ∈ [0,1]. Notice that *ϕ* = 1 means that this sign-pattern of the intersection line meets the requirements of sign-satisfaction for all the sample pairs. Therefore, the magnitude of the computed *ϕ* can be seen as the confidence of this potential solution to be a solution of the sign-satisfaction problem.
5. **Go back to Step (2) until Ψ > 1 intersection lines have been computed.**

In summary, selecting the intersection line can be seen as a “preference” sampling in the sign-satisfaction graph, because this intersection line can be crossed by at least (*N* − 1) hyperplanes in *M*_*i*_.

We illustrate the basic idea of the above discussion in the following example:

##### Example 7.

We compute the difference vector of all the sample pairs (the samples contains taxon 1) in Example 5 and stack them in the following matrix:

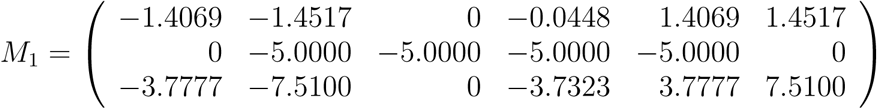

Each column of *M*_1_ is the difference of a sample pair, corresponding to the normal vector of a plane orthogonal to the associated (***x***^*I*^ − ***x***^*K*^). In Supplementary Fig. 5a, the intersection line (black line) is intersected by the planes where each of normal vectors respectively corresponds to the 1-st and 5-th column of the above *M*_1_. The black line crosses the regions with sign-pattern (−, 0, +)^⊤^ and (+, 0, −)^⊤^. At least these two regions have been crossed by two planes. Due to the fact that we know that *J*_11_ < 0, for the next step we need to count the number of the remaining hyperplanes that cross the region with the sign-pattern (−, 0, +)^⊤^. In Supplementary Fig. 5b we find that four of the remaining hyperplanes cross this intersection line, that is, the normalized *ϕ* satisfies *ϕ* =1. It means that the sign-pattern of this intersection line is the inference of `sign`(***J***_1_) because it meets the requirements of sign-satisfaction for all the sample pair.

#### 2.3.4. The heuristic algorithm combing sign-satisfaction and intersection of hyperplanes

Combining the sign-satisfaction graph with the sampling procedure described above, we propose a heuristic algorithm to infer the sign-pattern of ***J***_*i*_.

Our heuristic algorithm has two inputs: the steady-state dataset for the *i*-th taxon 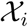 and a user-defined parameter Ψ determining how many intersection lines of hyperplanes will be constructed. The algorithm has of four steps, as described in Supplementary Fig. 6. Applying this procedure for *i* = 1,…,*N*, we can get the sign-pattern of the whole Jacobian matrix.

In summary, the algorithm works as follows. After generating an intersection line, we get the three orthants corresponding to this intersection line. Then we count how many hyperplanes cross the orthants determined by this intersection line using the sign-satisfaction graph, and this count can be normalized as *ϕ* ∈ [0,1] indicating the confidence of this potential solution to be a solution of final inference. Finally, we select the intersection line with the maximal *ϕ* among the generated Ψ intersection lines as the final inferred sign-pattern 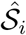.

Note that if the algorithm is stuck in generating an intersection line for some subset of (*N* − 1) hyperplanes, the heuristic algorithm will fail. Numerical experiments suggest this situation happens only when the data is not informative enough or the number of samples is smaller than the threshold Ω*. In Fig. 3 of the main text, we presents the results of the minimal number of samples Ω* required for a community with size *N*.

### 2.4. Limitations of the inference when using relative abundance data

High-throughput amplicon sequencing of 16S RNA has become a well-established approach for profiling microbial communities. The result of this procedure are measurements of the *relative abundance* of each taxa in the microbial community, meaning that these quantities have been normalized to sum to one (or some other arbitrary constant). This implies that an increase of the relative abundance of one taxon must be accompanied by a decrease in the relative abundance of other taxa. This severely limits the application of system identification methods based on temporal data, as discussed with details in [8, 9].

The use of steady-state samples containing relative abundance also leads to inference errors. Consider, for example, that there exists three relative abundance profiles containing taxon *i*, say 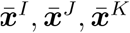. Since they are relative abundances, the sum of each of these samples must equal 1. This also implies that 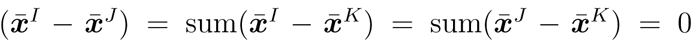. Consequently, the vector 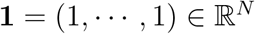 satisfies

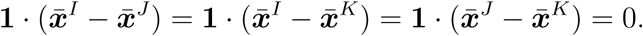

In other words, this vector 1 is always in orthogonal to all sample differences and the intersection line of the (*N* − 1) hyperplanes generated by relative abundance is always 1. Therefore, the heuristic algorithm fails in correctly inferring the sign-pattern of Jacobian matrix using relative abundances, because it always predicts that one possible sign-pattern is `sign`(1) = (+,…, +).

## 3. INFERRING THE TOPOLOGY OF ECOLOGICAL NETWORKS

In essence, inferring the zero-pattern is similar to inferring the sign-pattern. Indeed, it is only necessary to recognize any non-zero entry of the inferred sign-pattern as a non-zero entry in the inferred zero-pattern. Notice how the zero-pattern corresponds to hyperplanes exactly aligned to the orthants of 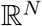. Therefore, any measurement noise will make the difference of sample pairs deviate from the axis, easily leading to inference errors (see Supplementary Fig. 1d,f). To alleviate this problem, we introduce a user-defined cutoff value to judge the zero-pattern of *J*_*ij*_ based on the angle between the axis and the intersection line (Supplementary Fig. 1).

For the brute-force method, first we set an element in the difference of sample pair ***x***^*I*^ − ***x***^*K*^ to 0 if the magnitude of that element is less than the user-defined cutoff. Second, we construct the hyperplanes respectively orthogonal to these modified difference of sample pairs. Third, we count how many hyperplanes cross each orthant in the 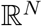. Finally, we select the region crossed by the maximal hyperplanes as the inferred zero-pattern. Recall that the brute-force method is limited to infer the microbial community with *N* ≤ 10.

For larger microbial communities, we also developed a heuristic algorithm that is very similar to our heuristic algorithm for inferring the sign-pattern. In that algorithm, notice how the deviation of an intersection line from an axis is directly given by its directional vector. Indeed, this vector contains the cosine of the angles between the axis and the intersection line. This algorithm works as follows:

1. **Construct the matrix of the difference of all the sample pairs.** Consider the set of all vectors 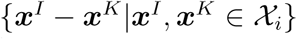. Let 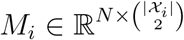 be a matrix constructed by stacking all the 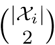 vectors, where 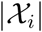 is the number of samples containing taxon *i*. By construction, each column of *M*_*i*_ is the normal vector of a hyperplane orthogonal to the difference of the corresponding sample pair.
2. **Randomly sample (*N* − 1) hyperplanes.** Choose randomly *N* − 1 columns from *M*_*i*_.
3. **Find the intersection of the (*N* − 1) sampled hyperplanes to obtain an intersection line.** This can be done by finding the kernel of the matrix obtained by stacking the chosen columns. Note that the randomly sampled (*N* − 1) hyperplanes not always intersect in a line, because some hyperplanes might be parallel. However, this situation is non-generic in 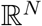. Thus, if the randomly sampled hyperplanes do no intersect as a line, we return to step 2 and Choose a new subset of columns.
4. Set the elements in the directional vector of this intersection line as zero if their absolute values are less than the cutoff value. Then we get a new directional vector. Note that we scale the 2-norm of this directional vector to 1. The absolute value of *i*-th element in the directional vector represents the cosine of the angle between the intersection line and *x*_*i*_-axis. If the value is large enough, it means the intersection line almost locates at the *x*_*i*_-axis. Therefore, if the absolute value of directional vector is smaller than the cutoff, we set this entry to 0.
5. **Count how many hyperplanes cross the region of the new directional vector using the sign-satisfaction graph.** The sign-pattern of new directional vector represent the orthants in 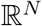 crossed by all those (*N* − 1) hyperplanes. For the rest hyperplanes of *M*_*i*_ (i.e., the rest of the columns in *M*_*i*_), let 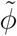 be the number of those hyperplanes that cross the orthants. We normalize 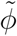 using 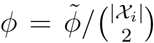, so that *ϕ* ∈ [0,1]. Notice that *ϕ* = 1 means that the sign-pattern of the new directional vector meets the requirements of sign-satisfaction for all the sample pairs. Therefore, the magnitude of the computed *ϕ* can be seen as the confidence of this potential solution to be a solution of the sign-satisfaction problem.
6. **Go back to Step (2) until Ψ ≥ 1 intersection lines have been computed.**

We validated this method using steady-state data generated from four different population dynamics. Except the Generalized Lotka-Volterra (GLV), the other three population dynamics models have non-linear functional responses: Holling Type II (H), DeAngelis-Beddington (DB) and Crowley-Martin (CM). Supplementary Fig. 7 shows the inferred network topology on four different population dynamics models. We found that in case the noise level is *η* = 0.1, the accuracy of inference can be around 0.8, if the cutoff is between 0.1 and 0.2. However, in the noiseless case, increasing the cutoff can decrease the accuracy, because larger cutoff induces more false positives of interactions.

## 4. INFERRING THE ECOLOGICAL INTERACTION TYPES

Using the brute-force method to infer the interaction types is deterministic because we search all the combinations of {+, 0, −}^*N*^, and accuracy increases with the increment of sample size. However, due to the time complexity, application of the brute-force method is limited to small microbial communities, e.g., *N* ≤ 10. This motivated us to develop the heuristic method of Supplementary Note 2 that is suitable for larger microbial communities.

To validate the effectiveness of our heuristic algorithm, we tested it using simulated steady-state data generated by models of the form Eq. (S1). In particular, we considered a model with pair-wise interactions of the form

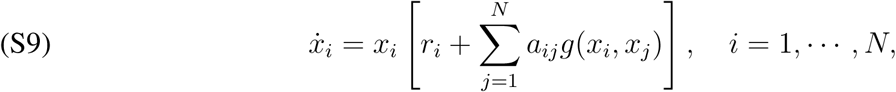

where 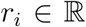 is the intrinsic growth rate of the *i*-th taxon, 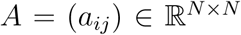 is a constant matrix and the function 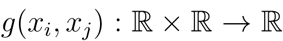 is the so-called *functional response* [1, 2, 3, 4, 5, 6]. Recall that these functional responses model the intake rate of a consumer as a function of food density, and thus different functional responses correspond to different mechanisms of interaction between taxa.

We used Eq. (S9) to generate synthetic steady-state datasets for 4 different functional responses with different complexity. The first was the linear functional response

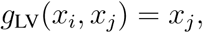

for which Eq. (S9) actually reduces to the classical Generalized Lotka-Volterra (GLV) model. In this case, the accuracy of the heuristic algorithm on inferring the sign-pattern `sign`(*J*) = `sign`(*A*) is 100% if there are enough steady-state samples, see Fig. 3a in the main text. Indeed, this is a consequence of the following proposition:

### Proposition 2.

In the noiseless case, if the functional response is linear, the directional vector of intersection line of any (*N* − 1) hyperplanes orthogonal to 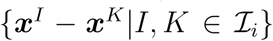 is the same and parallel to ***J***_*i*_.

*Proof.* Due to the fact that the functional response is linear, the Jacobian matrix become simple and constant for different samples, that is, *J = A*. Therefore, Eq. (S3) is equal to

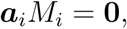

where 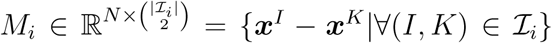 denotes the difference of all sample pairs. As we know, ***a***_*i*_ ≠ 0, representing the interaction vector in the A matrix, is unique to *M*_*i*_. Thus the non-trivial solution of ***a***_*i*_ in the above equation array must meet the requirement

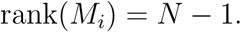

That is to say, if we randomly select (*N* − 1) columns in *M*_*i*_ as 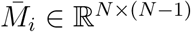, then

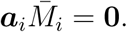

Actually, the randomly selected 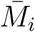 corresponds to (*N* − 1) hyperplanes respectively orthogonal to each columns of 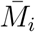 in the geometric perspective. The directional vector of intersection line of these (*N* − 1) hyperplanes can be calculated by null 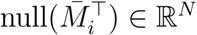, which is parallel to ***a***_*i*_.

The remaining three functional response were Holling Type II (H), DeAngelis-Beddington (DB) and Crowley-Martin (CM), given by the following equations

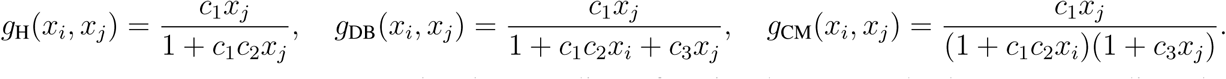

Here *c*_1_, *c*_2_, *c*_3_ are constants. Note that these nonlinear functiunal responses lead to more complicated population dynamics. For the results presented in Fig. 3 of the main text, we used *c*_1_ = 1, *c*_2_ = *c*_3_ = 0.1. Thuse results show that the heuristic algorithm accurately infers the sign-pattern of Jacobian matrix for these three functional responses and its accuracy is above 95%.

## 5. INFERRING INTERACTION STRENGTHS WITH GLV DYNAMICS

A particular class of systems in (S1) is when the Jacobian *J*_*i*_ is constant, implying that 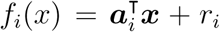 for some constant vector 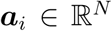 and scalar *r*_*i*_. In such case, the system reduces to the Generalized Lotka-Volterra (GLV) model

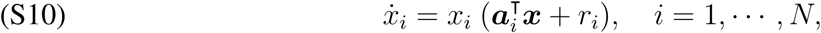

where 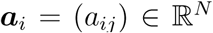 is the *i*-th row of the so-called interaction matrix 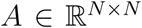, and 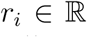 is the intrinsic growth rate of taxon *i*. As discussed in the main text, the GLV models also allows defining the *interaction strength* of taxon *j* on taxon *i* as *a*_*ij*_.

### 5.1. A condition for detecting GLV dynamics

Our first observation is that the steady-state samples 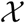 can be used to decide if they could be produced by a GLV model:

#### Theurem 3.

A necessary condition for the dynamics of the *i*-th species to be GLV is that all samples 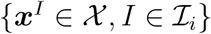 align into a hyperplane.

*Proof.* If for all 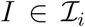 the samples *x*^*I*^ align into a hyperplane, then *f*_*i*_(*x*) should be a hyperplane whose general equation is 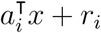.

As discussed in the main text, with real data containing measurement noises and other errors, the samples will not align exactly into a hyperplane. In such case, the coefficient of determination (denoted by *R^2^)* of a hyperplane fitted to the samples containing taxon *i* can be used to judge if its dynamics can be adequately described by the GLV model. For a given dataset 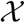, if the average of *R*^2^ of the hyperplanes fitted to the samples of the *i*-th taxon is > 0.9, then we consider that it is possible to infer the inter-taxa interaction strengths and intrinsic growth rates using the GLV model for this taxon. Otherwise, we recommend to infer only the interaction types. The pipeline for detecting GLV dynamics is described as Supplementary Fig. 8.

### 5.2. Inference of interaction strengths and intrinsic growth rates

Under the GLV model, Eq. (S3) reduces to

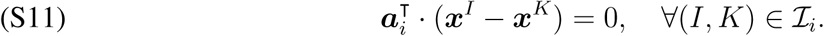

If we denote by *P*_*i*_ the (*N* − 1) dimensional hyperplane spanned by all the steady-state samples sharing the *i*-th taxon 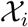, Eq. (S11) implies that the ***a***_*i*_ belongs to the one-dimensional space orthogonal to *P*_*i*_. Thus the normal vector of the fitted hyperplane according to 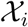 is parallel to ***a***_*i*_. To infer the precise value of interaction strengths, additional prior information, at least one non-zero element in ***a***_*i*_, is needed. Otherwise, we can only infer the relative strength of the interactions between taxa.

### 5.3. Applying the Knockoff filter to control the false discovery rate

Eq. (S11) shows that ***a***_*i*_ can be inferred by fitting a hyperplane based on all the steady-state samples sharing the *i*-th taxon 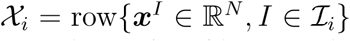, provided that we know at least one non-zero element in ***a***_*i*_, say *a*_*ii*_ (or an estimate 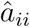 of it). Consider that the ecological network to be inferred is sparse. Then, a natural method to find a sparse solution is by using the so-called Lasso regression:

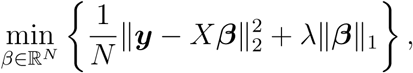

where λ is the Lasso (regularization) parameter. Here ***y*** is the *i*-th column of 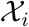, and

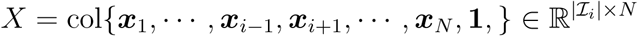

is the matrix obtained from 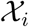 by deleting the *i*-th column and adding **1** in the end. ***x***_*i*_ is the *i*-th column of 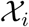. This structure happens because for the GLV we have *r*_*i*_ + *a*_*i*1_*x*_1_ +…+ *a*_*ii*_*x*_*i*_ + *a*_*i*_,_*i*+1_*x*_*i*+1_ +…+ *a*_*iN*_*x*_*N*_ = 0 and we assumed for the numerical results that *a*_*ii*_ = −1. Once a solution *β* to the above Lasso problem is found, the estimation 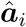 for ***a***_*i*_ is given by

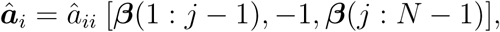

where ***β***(*i*_0_: *i*_*f*_) is the vector obtained by concatenating the elements *i*_0_ to *i*_*n*_ of the vector ***β***. Recall that the parameter *λ* in the Lasso is crucial for accuracy. A classical method to optimally choose this parameter is using cross validation.

However, even after using cross validation, the Lasso tends to induce a high false discovery rate (FDR), i.e., many zero interactions are inferred as non-zeros ones. Formally, the FDR of a inference procedure ***y*** = *X**β*** + ***z***, returning the inferred parameters 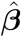, is defined as

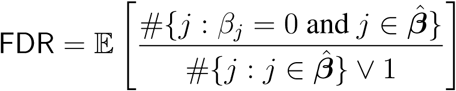

Here *a* ⋁ *b* = max{*a, b*}.

Recently, the so-called Knockoff filter has been proposed as an enhancement to the Lasso algorithm to maintain the FDR below a certain user-defined level *q* > 0, regardless of the value of the coefficients ***β*** (see [10]). This method works by constructing the so-called “knockoff variables” that mimic the correlation structure found in the real data. The knockoff copy of each variable act as a “control group”, allowing to assign a “trust” to each inferred variable. It has been shown this strategy successfully controls the FDR. In our work we used the Matlab package of the Knockoff filter as provided in https://web.stanford.edu/~candes/Knockoffs/package_matlab.html. The validation of the network inference with GLV dynamics is shown in Fig. 4 of the main text.

### 5.4. Blinded inference of interaction strengths by assuming GLV dynamics

Here we show that, if the steady-state samples were collected from a microbial community without GLV dynamics, the inference of interaction strengths by assuming GLV dynamics systematically leads to inference errors.

To illustrate this point, we first generated steady-state samples using Holling Type-II functional response. Then, we applied the GLV-based inference method to the steady-state samples in order to infer the interaction strengths. Supplementary Fig. 9 shows that the accuracy (the percentage of correct sign of the inferred interaction strengths compared with the sign of ground truth) of inferred results is very low, even in the absence of noise. This is consistent with the small value of *R*^2^ of fitted hyperplanes, which describe the deviations of samples to those fitted hyperplanes. This suggest that inferring the interactions strengths of a real microbial community without first testing if its dynamics can be described by the GLV model can produce significative errors.

## 6. REAL DATASETS

### 6.1. A synthetic microbial community of 8 soil bacteria

In [11] a set of eight heterotrophic soil-dwelling bacterial species were studied for predicting species persistence in different assembled microbial microcosms. The steady-state dataset 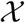 consists of a total of 101 different species combinations: 8 solos, 28 duos, 56 trios, 8 septets and 1 octet (Supplementary Fig. 10a). Each species combination of cultivation was carried out in duplicate and started from different configurations of initial abundance. We averaged the steady states from different initial conditions.

First we find that *R*^2^ of each fitted hyperplane is less than 0.9 (Supplementary Fig. 10b), which indicates that this microbial community could not be properly described by the GLV model. Hence we focus on the inference of interactions types between any two species. To be fair, without considering the 8 solos and 28 duos, we analyze the rest steady-state samples. We use both the brute-force algorithm (see Fig. 5 of main text) and the heuristic algorithm (Supplementary Fig. 10c,d) to infer the ecological interaction types. In Supplementary Fig. 10c, blue (or red) means inhibition (or promotion) effect of species *j* on species *i*, respectively. We found that 11 signs were falsely inferred, 5 signs were undetermined by the analyzed steady-state samples. The inferred results are very similar with the brute-force method shown in Fig. 5b of main text. Furthermore, Supplementary Fig. 10d shows that once Ψ is larger than a certain value, the accuracy in the inference does not increase any more.

### 6.2. A synthetic community of maize roots with 7 bacterial species

(Fig. 6b in the main text). There are in total 7 bacterial species (Ecl, Sma, Cpu, Opi, Ppu, Hfr and Cin) in this community [12]. The available steady-state data consists of 7 sextets (i.e., data from seven experiments in which six different species grow together) and 1 septet (i.e., data from one experiment in which the seven species grow together). This leads to a total of 8 steady-state samples, see Supplementary Fig. 11a.

First, based on our theoretical result showing that in the generalized Lotka-Volterra (GLV) model the steady states that share common species will align into a hyperplane, we concluded that this bacterial community does not follow the GLV dynamics (see Supplementary Fig. 11b). Thus, we have to focus on inferring the interaction types, rather than interaction strengths.

Second, only using the 7 sextets we inferred the sign-pattern of the Jacobian matrix (Fig. 6a in the main text). Based on the inferred sign of *J*_*ij*_, we can predict how the abundance of species *i* will change, when we add species *j* to the community (see results in the Main Text).

### 6.3. A synthetic microbial community of two cross-feeding partners

In this community [13], two non-mating strains of the budding yeast, *Saccharomyces cerevisiae,* were engineered to be deficient in the biosynthesis of one of two essential amino acid tryptophan (Trp) or leucine (Leu), and to overproduce the amino acid required by their partner. It has been demonstrated that these two strains form a community with cross-feeding mutualism, where each strain provides the amino acid needed by its partner. In [13], the authors inoculated monocultures and co-cultures at a range of concentrations of supplemented amino acids in a well-mixed liquid batch. Supplementary Fig. 12a-c shows the abundance of the co-cultures and monocultures for the Trp and Leu strains at low, medium and high levels of supplemented amino acids. After 7 days cultivation, the abundance of each species approaches its steady state. Note that for each scenario, the experiments inoculate a constant amount of resources at the beginning. Here the type of interaction is defined by comparing the abundance of co-cultures with monocultures at the end of cultivation. As the supply of amino acids increases from low, to medium to high concentrations, the interaction between this pair of strains shifts from obligatory mutualism (Supplementary Fig. 12a), to facultative mutualism (Supplementary Fig. 12b), and to parasitism (Supplementary Fig. 12c), respectively.

We applied our inference method to each scenario. Supplementary Fig. 12d-f shows the diagrams of our inference results that are consistent with the empirical observations. For example, in Supplementary Fig. 12e,f, the cyan line orthogonal to the red line is very close to the Leu axis, which indicate the effect of Trp on Leu is very weak. Especially in Supplementary Fig. 12f, this promotion effect can be ignored.

### 6.4. A synthetic community of 14 auxotrophic *Escherichia coli* strains

Starting from a prototrophic E. coli derivative MG1655, the authors of [14] generated 14 strains, each containing a gene knockout that lead to an auxotrophic phenotype unable to produce 1 of 14 essential amino acids. By convention, the authors labeled each auxotrophic strain by the amino acid it lacks. For example, the methionine auxotroph Δ*metA* auxotroph is strain M. It was confirmed that the 14 auxotrophs *(C, F, G, H, I, K, L, M, P, R, S, T, W, Y)* show no growth in M9-glucose minimal media after 4 days. Indeed, they grow only when supplemented with the essential amino acid they were not able to produce. This dataset consists of co-cultures of all 91 possible strain pairs from the 14 characterized auxotrophic strains. For each pairwise co-culture, we are able to calculate the total fold growth, i.e., the yield of the community calculated by (total final cell density)/(total initial cell density), as well as the fold growth of each strain. Since these auxotrophic strains cannot grow by themselves, if strain *i* is able to grow as a co-culture when paired with strain *j*, and strain *i*’s fold growth is *F*_*ij*_ > 1, this implies that strain *j* promotes the growth of strain *i,* i.e., *J*_*ij*_ > 0. By contrast, if *F*_*ij*_ < 1, we cannot conclusively say that *J*_*ij*_ < 0 because we lack the monoculture data. Therefore, the fold-growth metric can only be used to detect a promotion effect between two strains.

First, we found that *R*^2^ of all fitted hyperplanes are smaller than 0.9, implying that the population dynamics of this microbial community cannot be properly described by the GLV model (Supplementary Fig. 13a). Second, we used the heuristic algorithm to infer the interaction types (Supplementary Fig. 13b). Note that the complexity of the inference approaches 3^14^ ~ 4 x 10^6^ if we use the brute-force algorithm. We found that the types of 14 pairwise interactions cannot be determined with the given dataset (marked in gray in Supplementary Fig. 13b). Third, we showed the fold growth matrix *F* = (*F*_*ij*_) from experimental observations (Supplementary Fig. 13c), with *F*_*ij*_ the fold growth of strain *i* (row) in the co-culture paired with strain *j* (column). Here we set *F*_*ij*_ ≥ 20 as an indication of promotion effect of strain *j* on strain *i*. There are in total 71 promotion interactions with such a large confidence (shown in red, Supplementary Fig. 13c). We will use them as the ground truth to check our inference results on promotion effects (i.e., positive signs, shown in red in Supplementary Fig. 13b). We found we inferred 13 wrong positive signs (marked as ‘×’ in Supplementary Fig. 13c), and missed 5 positive signs (marked as ‘?’ in Supplementary Fig. 13c). Therefore, our inference of positive signs has an accuracy of 74.65% (53/71), if we set the fold growth threshold 20 as the indication of promotion effect. We also observed that the accuracy on the inference generally increased by increasing this threshold (Supplementary Fig. 13d).

## 7. RELATIONSHIP TO EXISTING NOTIONS OF INTER-TAXA INTERACTIONS

In Assumption 2, we considered that the Jacobian of (S1) determines the interaction types between microbial taxa. This assumption was then used to build our network reconstruction method. Here we discuss how this consideration compares to other existing definitions and notions of “interactions” available in the ecological literature.

In general ecological systems, understanding the interactions between taxa and their strengths is key for developing predictive models and conservation strategies. This has motivated the introduction of several empirical indices for inter-taxa interactions, specially for consumer-prey ecosystems [15, 16]. Let *x*_1_ and *x*_2_ denote the abundances of prey and consumer, respectively. Consider two samples for this ecosystem consisting of an experiment with the prey in isolation 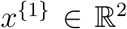 —that is, with the consumer or predator deleted— and other with both prey and consumer present 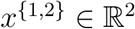. Here we discuss the four empirical indices as used in [16]:

a. Raw difference: 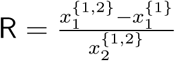
b. Paine’s index: 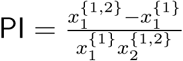
c. Community importance: 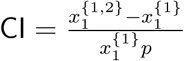, where 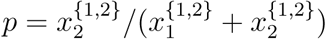
d. Dynamic index: 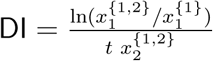, where *t* is time.

### Remark 7.

a. All the above indices have identical signs, solely determined by 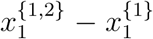. As shown in Example 3, such sign coincides with one of the possible sign-patterns obtained by applying our reconstruction method for *N* = 2 taxa. More precisely, the above indices coincides with our reconstruction method provided we assume this as prior information. Such prior information can be interpreted as adopting a “convention” for the sign of self-interactions (i.e., a kind of “relative sign-pattern”).
b. Compared to the analysis in [16], our reconstruction method provides more general conditions under which the above indices provide the correct sign of the interactions according to a mathematical model.
c. Our reconstruction method also generalizes the application of the above indices to ecosystems with an arbitrary number of taxa, and beyond the consumer-prey interactions.
d. Our reconstruction method provides conditions under which the available steady-state data is informative enough to infer the correct sign of a desired microbial interaction.
e. According to our framework, note there are two different interactions that is possible to infer: *x*_1_ → *x*_2_ and *x*_2_ → *x*_1_. The above indices and discussions are concerning the interaction *x*_2_ → *x*_1_ —that is, the effect of the consumer on the prey. In order to infer the sign of the interaction *x*_1_ → *x*_2_, we need to evaluate 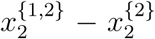. In the case of consumer-prey ecosystem with *N* = 2 taxa, it might be impossible to measure a non-zero ***x***^{2}^, since it corresponds to a steady-state abundance of consumers in the absence of prey. In such case, the set 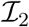 contains only one sample ***x***^{1,2}^, and thus the given data is not informative enough to infer this interaction. This argument could explain cases when for *N* taxa it is impossible to infer some interaction due to the absence of the needed sample, simply because in the absence of some taxa other become extinct.

---

**SUPPLEMENTARY FIGURE 1.**
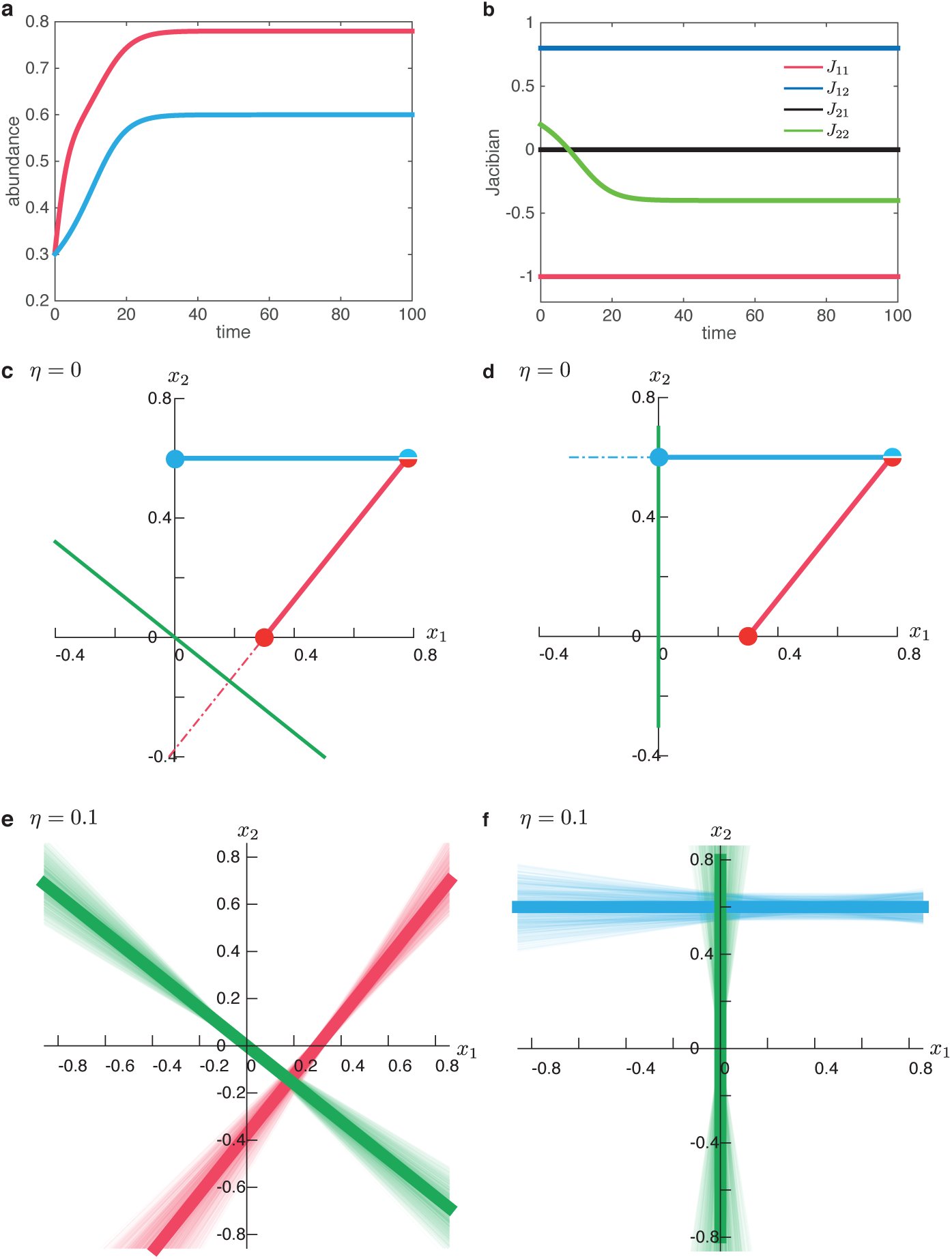
Inferring the zero-pattern of the Jacobian matrix. **a**. The temporal evolution of the abundance of each species. **b**. The sign of *J*_22_ is time varying, while the signs of the other elements in the Jacobian matrix are time-invariant. **c.** Inference of *J*_12_. According to the position of ***x***^{1,2}^ − ***x***^{1}^, the green line which is orthogonal to the red line cannot produce a zero entry for *J*_1_, implying that *J*_11_ ≠ 0 and *J*_12_ ≠ 0. This is consistent with the ground truth. **d**. Inference of J_12_. The green line (orthogonal to the blue line) is aligned with the *x*_2_-axis, indicating that *J*_21_ = 0 and *J*_*22*_ ≠ 0, consistent with the ground truth. **e**. When noise level *η* = 0.1, the light green line is orthogonal to the light red line corresponding to the difference of two noisy samples ***x***^{1}^ and ***x***^{1,2}^. The bold red and green lines correspond to the noiseless case. There are in total 1000 different measurements (replicates). We found that the angles between the green lines and *x*_1_-axis are large enough, letting us conclude that J_12_ ≠ 0 even if there exists some noise. f. When noise level *η* = 0.1, the light green line is orthogonal to the light blue line corresponding to the difference of two noisy samples ***x***^{1}^ and ***x***^{1,2}^. The bold blue and green lines correspond to the noiseless case. There are in total 1000 replicates. Among the 1000 replicates, the light green line is equally distributed to the left and right side of *x*_*2*_-axis, indicating that the deviation of the light green line from the *x*_2_-axis is likely due to measurement noises. This behavior let us introduce a user-defined cutoff value to judge the zero-pattern of *J*_*ij*_ based on the angle between the *x*_*1*_-axis (or *x*_*2*_-axis) and green lines.

**SUPPLEMENTARY FIGURE 2.**
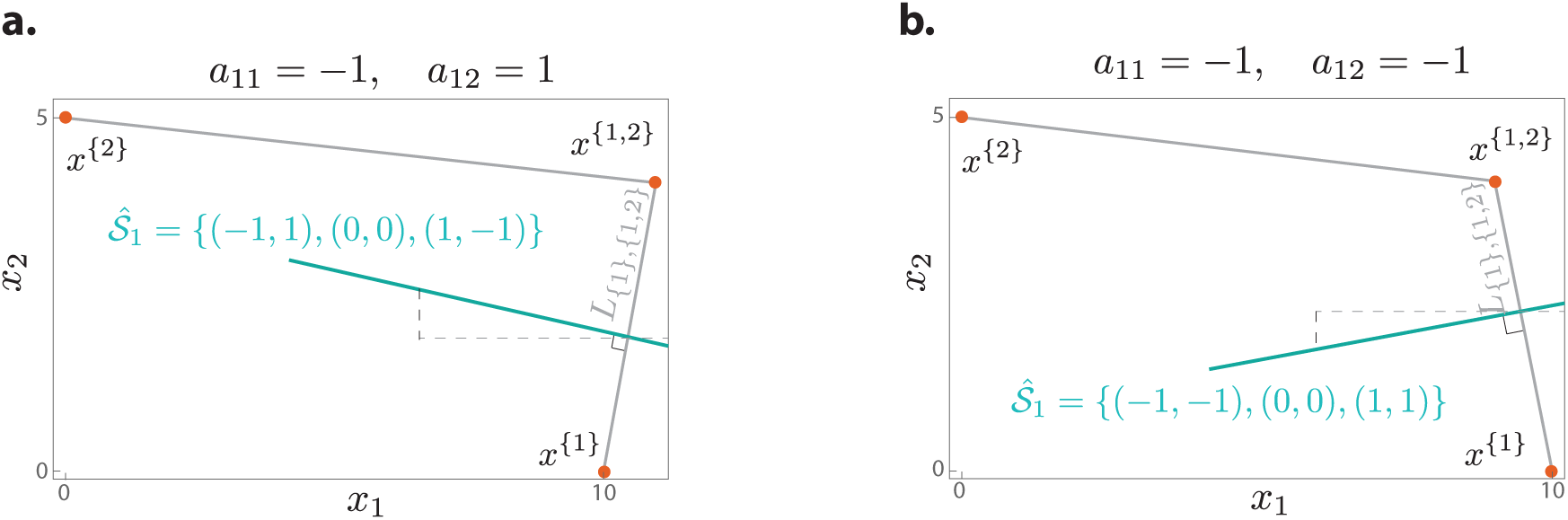
Reconstructing `sign`(***J***_1_) for *N* = 2 in Example 3.

**SUPPLEMENTARY FIGURE 3.**
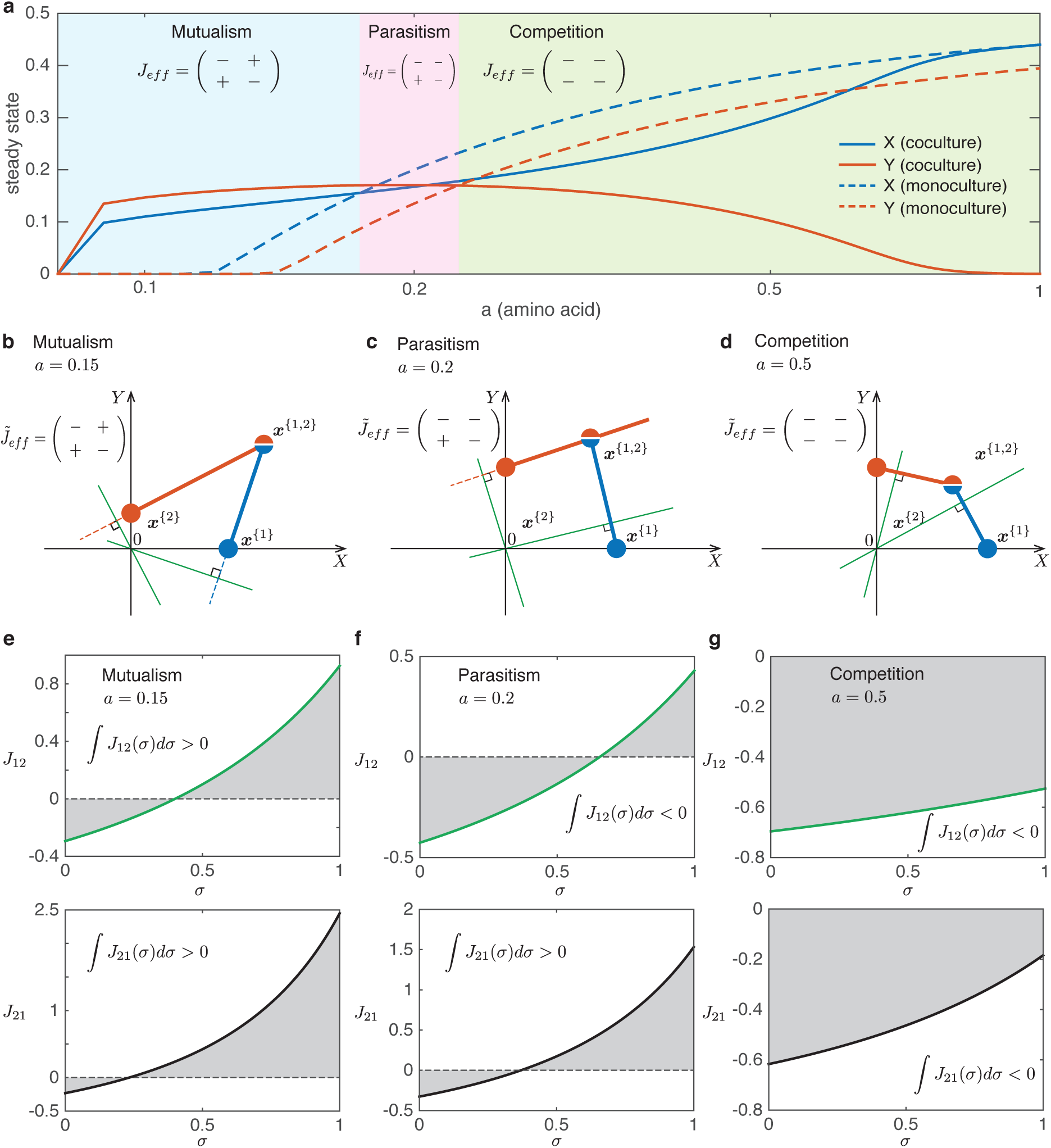
In case the sign-pattern of the Jacobian matrix is time varying, the results of our inference can be interpreted as the overall inhibition or promotion impact between different taxa. Here we consider a toy model of two species *X* and *Y*. Each has a per capita growth rate that is modulated by its mutualistic partner as well as the resource amount (denoted as *a*). The population dynamics model is shown in Eq. (S6), with model parameters *κ* = 0.12, *δ* = 0.5, *β* = 2. **a.** Three regimes of the interaction types emerge from different resource amount, from mutualism, parasitism to competition. The ground truth of the interaction types is determined by comparing the abundance of coculture (solid lines) with that of monoculture (dashed lines). **b-d.** Diagrams of our inference method under different resource amount. **e-g.** *J*_*ij*_(***x***^*I*^ + *σ* (***x***^*K*^ − ***x***^*I*^)) as a function of *σ* under different resource amount.

**SUPPLEMENTARY FIGURE 4.**
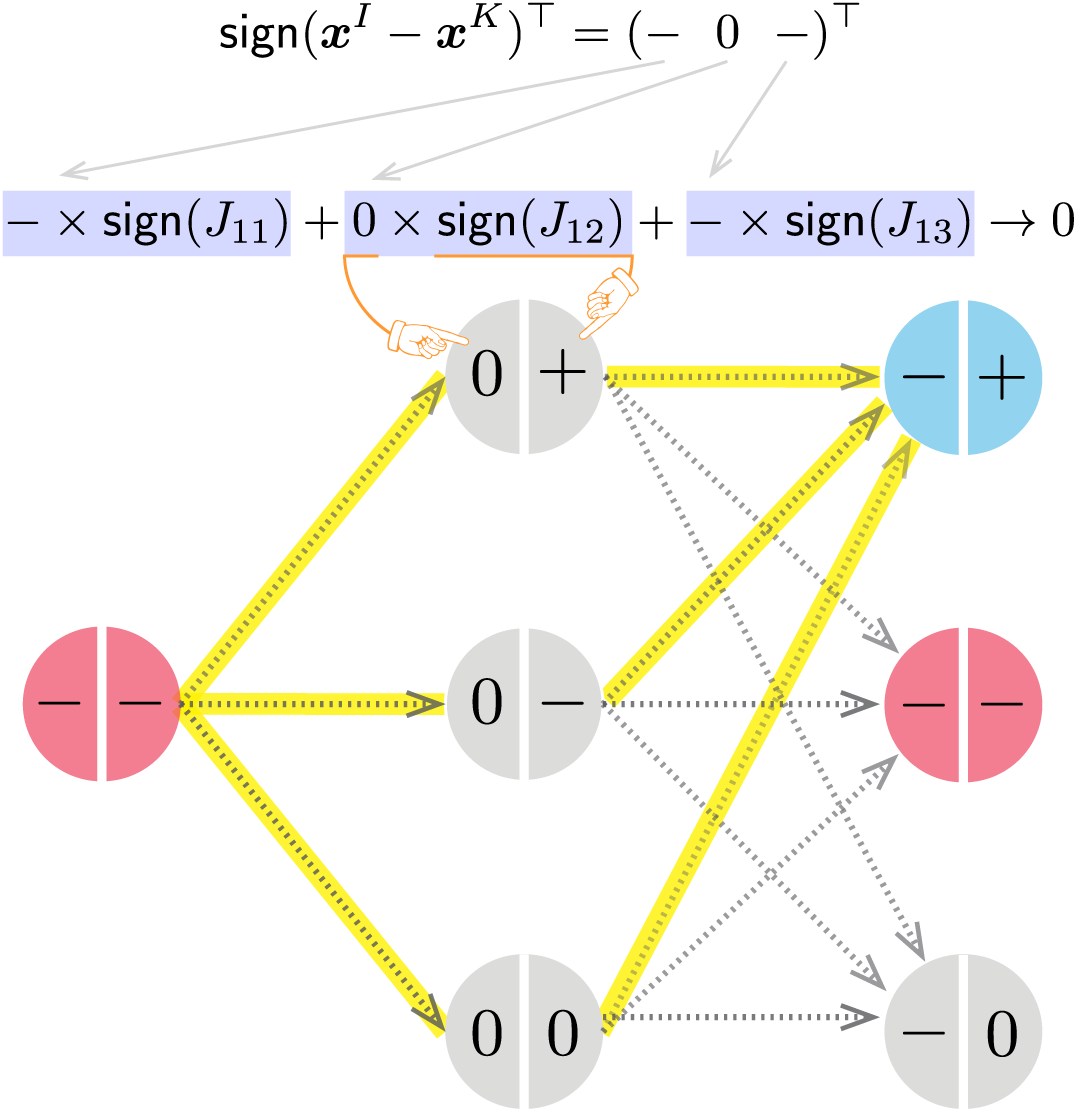
Sign-satisfaction graph and its solution for Example. 6. Each node is divided in two parts: the left is an entry of `sign`(***x***^*I*^−***x***^*K*^) and the right is an entry of `sign`(*J*_1_). The color of each node represents the multiplication of the sign of left and right part (red is positive, gray is zero and blue is negative). The yellow paths are the solutions to the sign-satisfaction graph, determining sign(*J*_1_), i.e., (−, +, +), (−, −, +), and (−, 0, +). Note that since we assume *J*_11_ < 0 as prior information, the first column of the sign-satisfaction graph only has one node.

**SUPPLEMENTARY FIGURE 5.**
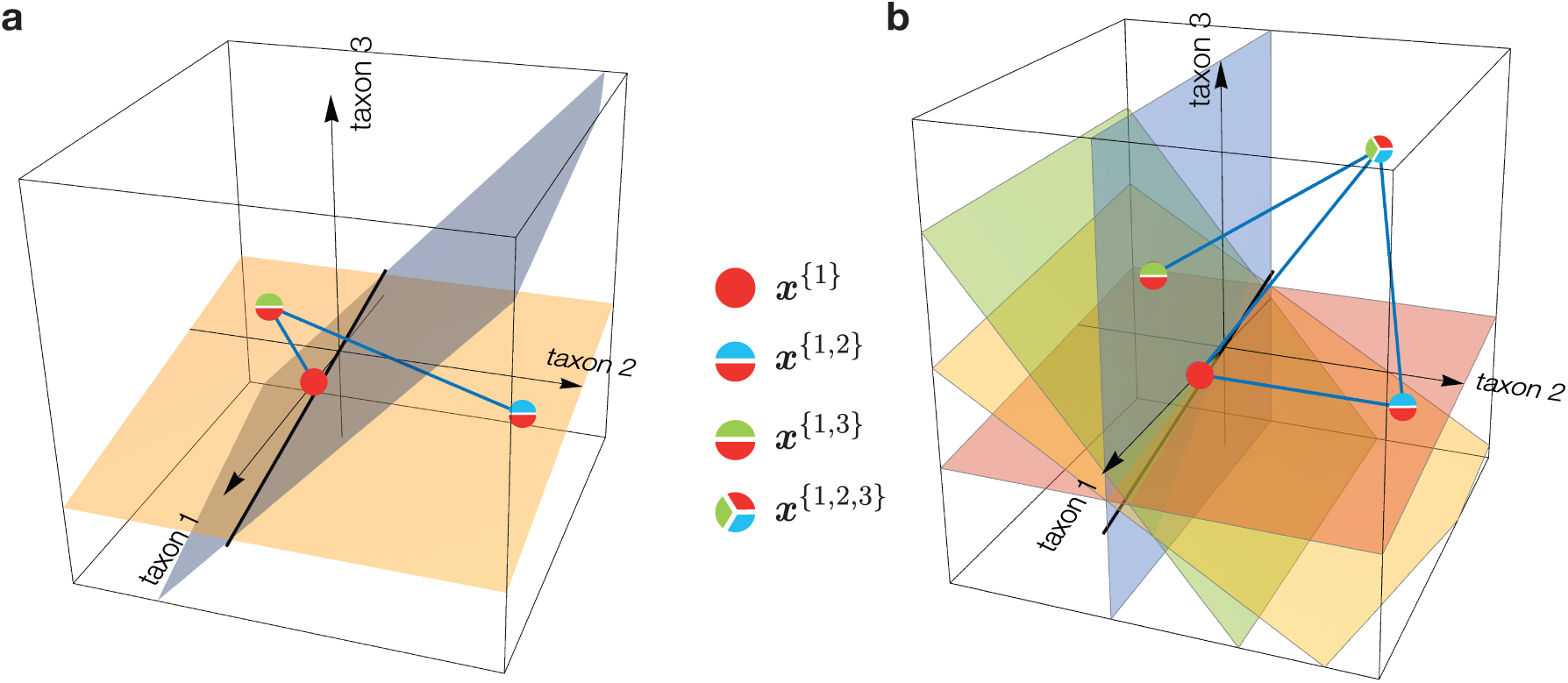
Intersections of planes provides a preference sampling in the sign-satisfaction graph. **a.** The black line is the intersection of the light orange and blue planes. Each of normal vectors (blue lines) corresponds to the first and fifth column of the matrix *M*_*1*_ in Example 5, respectively. Note that the orthants to which the intersection line belongs implies that those orthants are at least crossed by two planes. **b.** If there 4 samples sharing taxon 1, we will have a total of 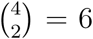 planes. Thus, *ϕ* will be the normalized count of how many of those hyperplane cross the orthants determined by the black intersection line.

**SUPPLEMENTARY FIGURE 6.**
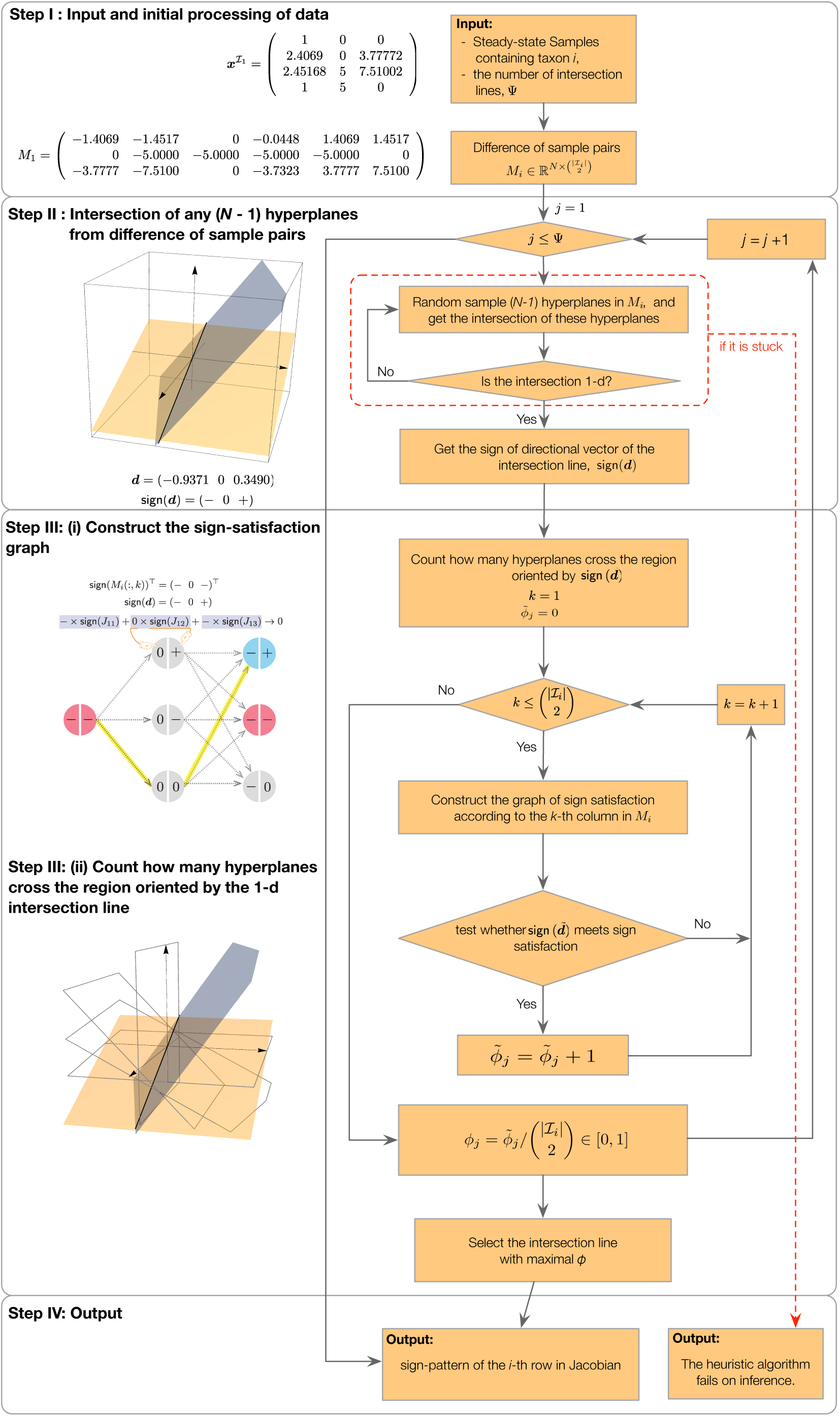
Pipeline of the heuristic algorithm for inferring interaction types.

**SUPPLEMENTARY FIGURE 7.**
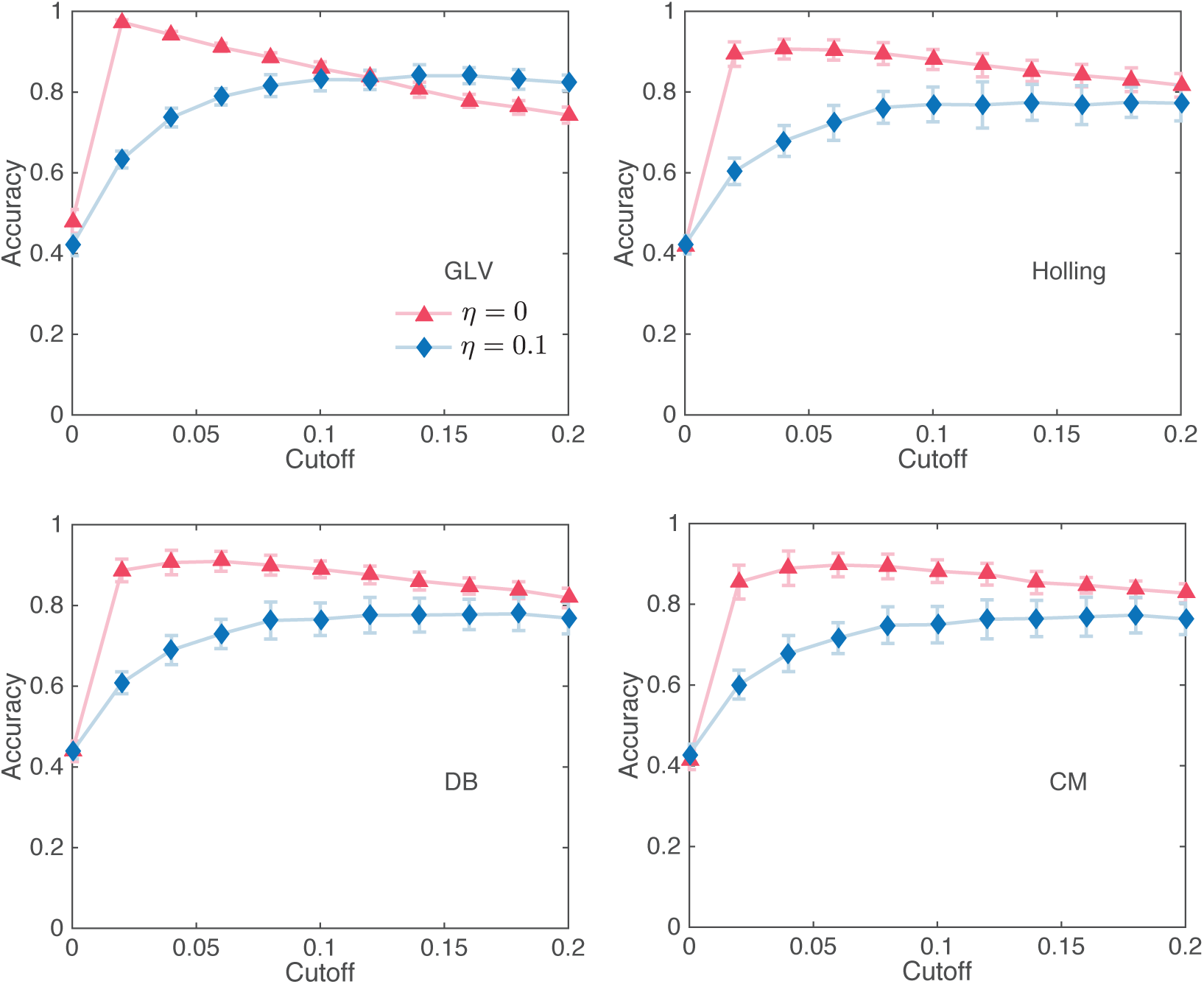
The inference of network topology for four different population dynamics models. Here the ecological network is generated using random network with *N* = 20 taxa and the connectivity equal to 0.4. The simulated steady-state samples are generated using the constants *c*_1_ = 1, *c*_2_ = *c*_3_ = 0.1 for Holling, DB and CM. The inference used Ω = 5*N* = 100 samples and Ψ = 5*N* = 100 intersection lines in the heuristic algorithm. The error bars represent standard deviation for 10 different realizations.

**SUPPLEMENTARY FIGURE 8.**
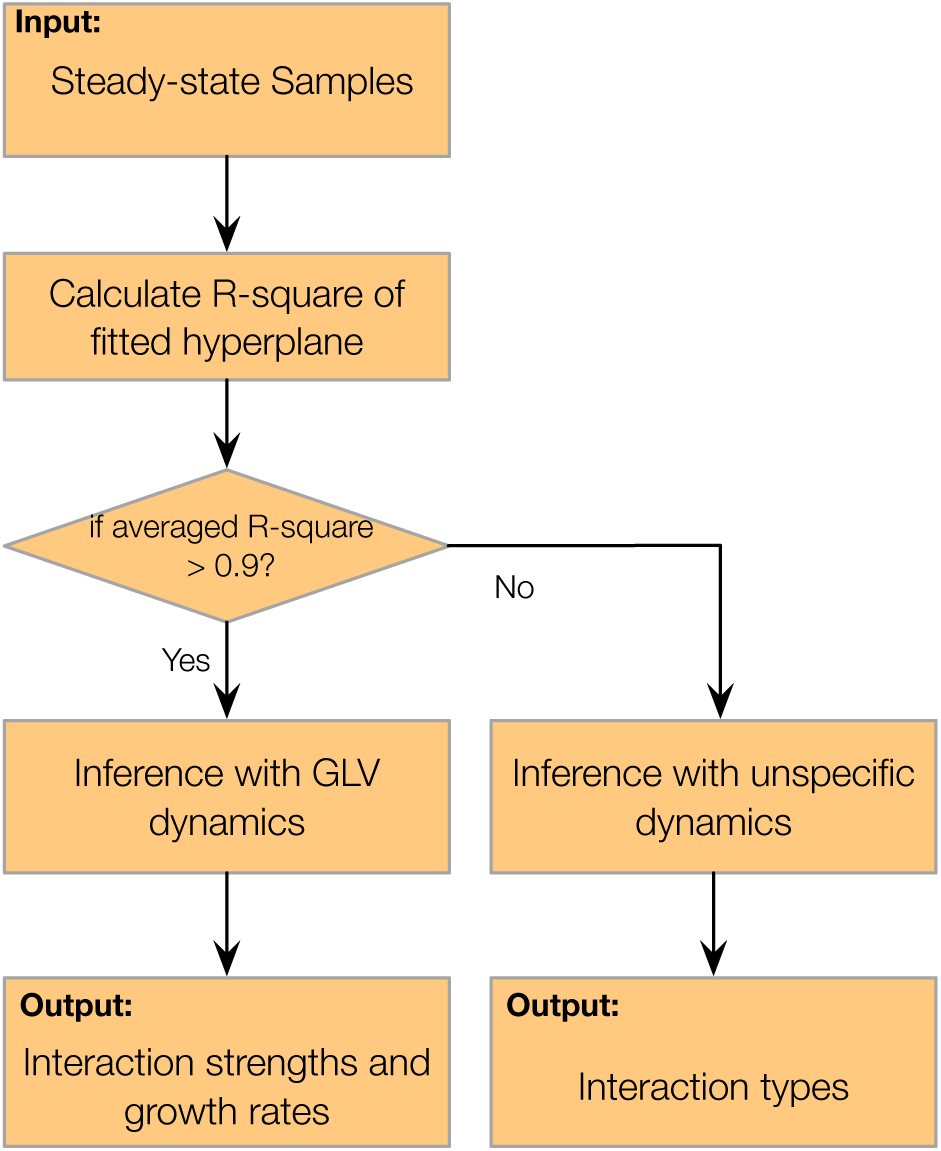
Pipeline of the heuristic algorithm for inferring interaction types.

**SUPPLEMENTARY FIGURE 9.**
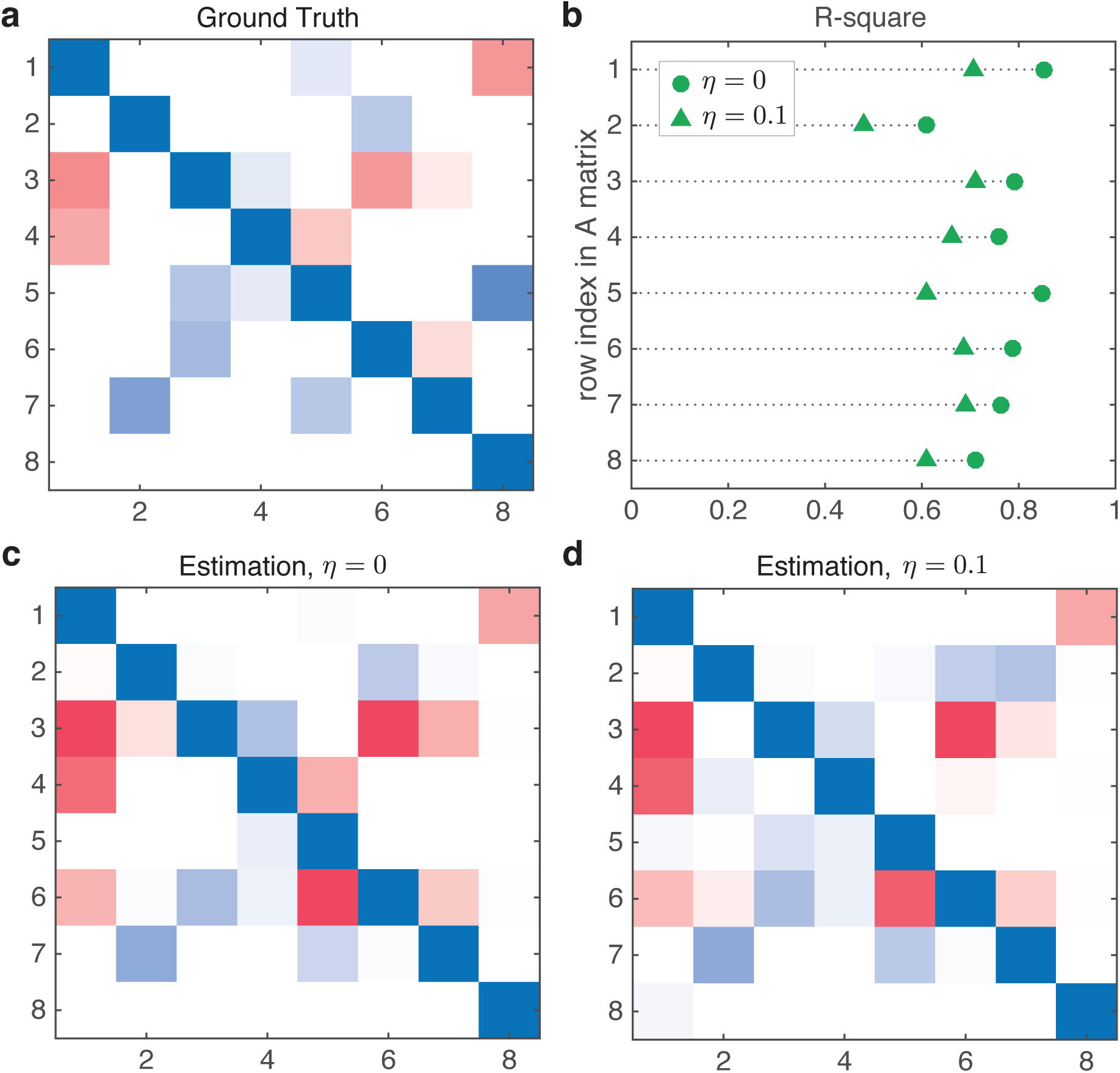
Inference assuming GLV dynamics for a microbial community without GLV dynamics. **a.** Here we generated the steady states from a microbial community of *N* = 8 taxa with Holling Type-II functional response. The A matrix is shown here. **b.** *R*^2^ of fitted hyperplanes in the noiseless (circle) or noisy (triangle) samples. **c,d.** The inferred A matrix. The accuracy of the inference in the noiseless and noisy cases are 0.7812 and 0.6719, respectively.

**SUPPLEMENTARY FIGURE 10.**
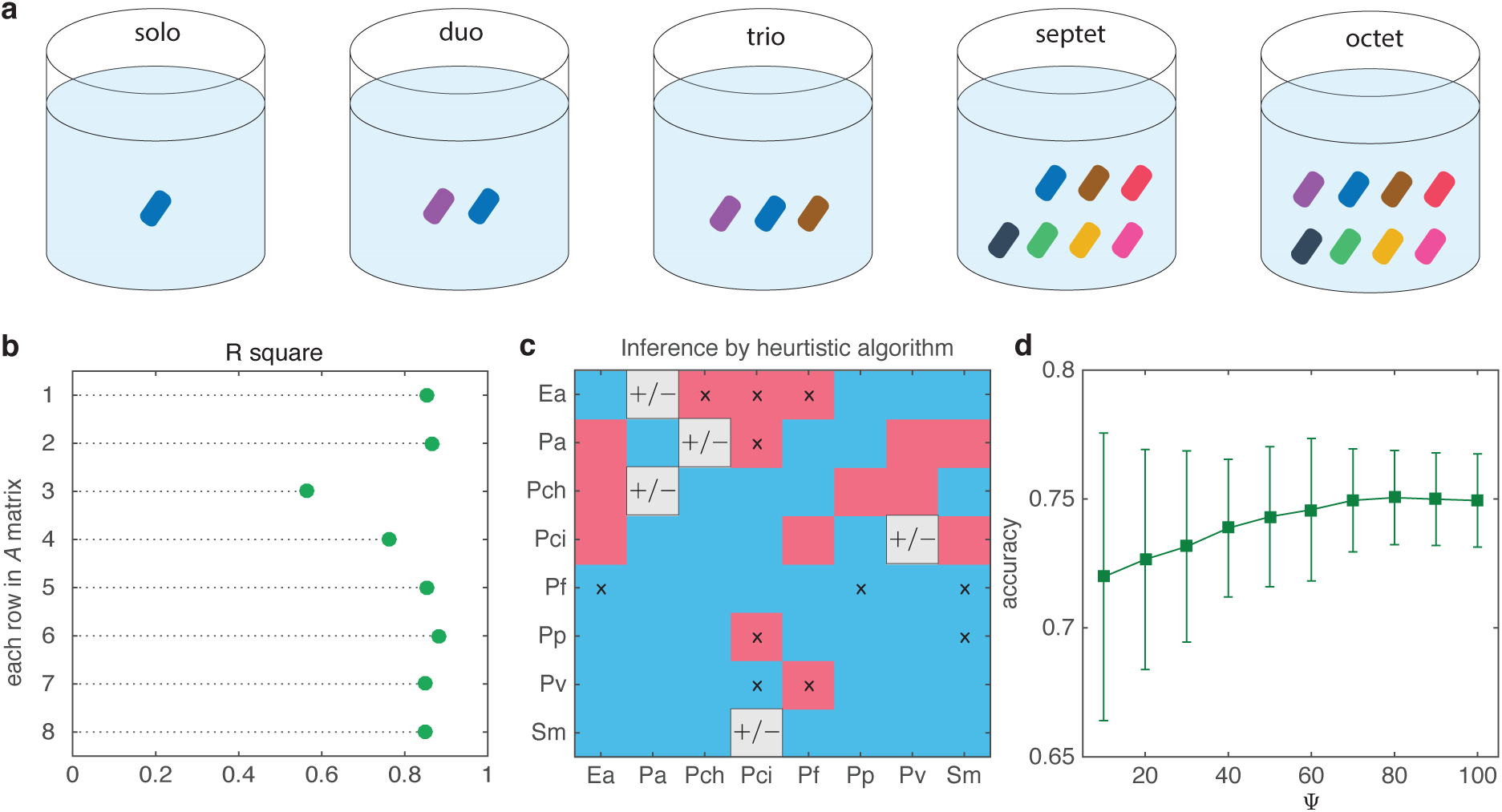
Inference of interaction types of a synthetic soil microbial community using our heuristic algorithm. **a.** 101 different species combinations: all 8 solos, 28 duos, 56 trios, all 8 septets, and 1 octet. **b.** We find that *R*^2^ of all fitted hyperplanes are smaller than 0.9. This suggests that the given samples cannot be properly described by the GLV model, and we should focus on the inference of interaction types. **c.** Inferred interaction types. Compared with the ground truth in Fig. 5a in the main text, there are 11 falsely inferred signs (false labels) and 5 signs cannot be determined by the given samples. We take the number of intersection line as Ψ = 50. **d.** The accuracy as a function of Ψ. Once Ψ is larger than a certain value, the accuracy could not increase any more. The error bar represents standard deviation for 30 different realizations.

**SUPPLEMENTARY FIGURE 11.**
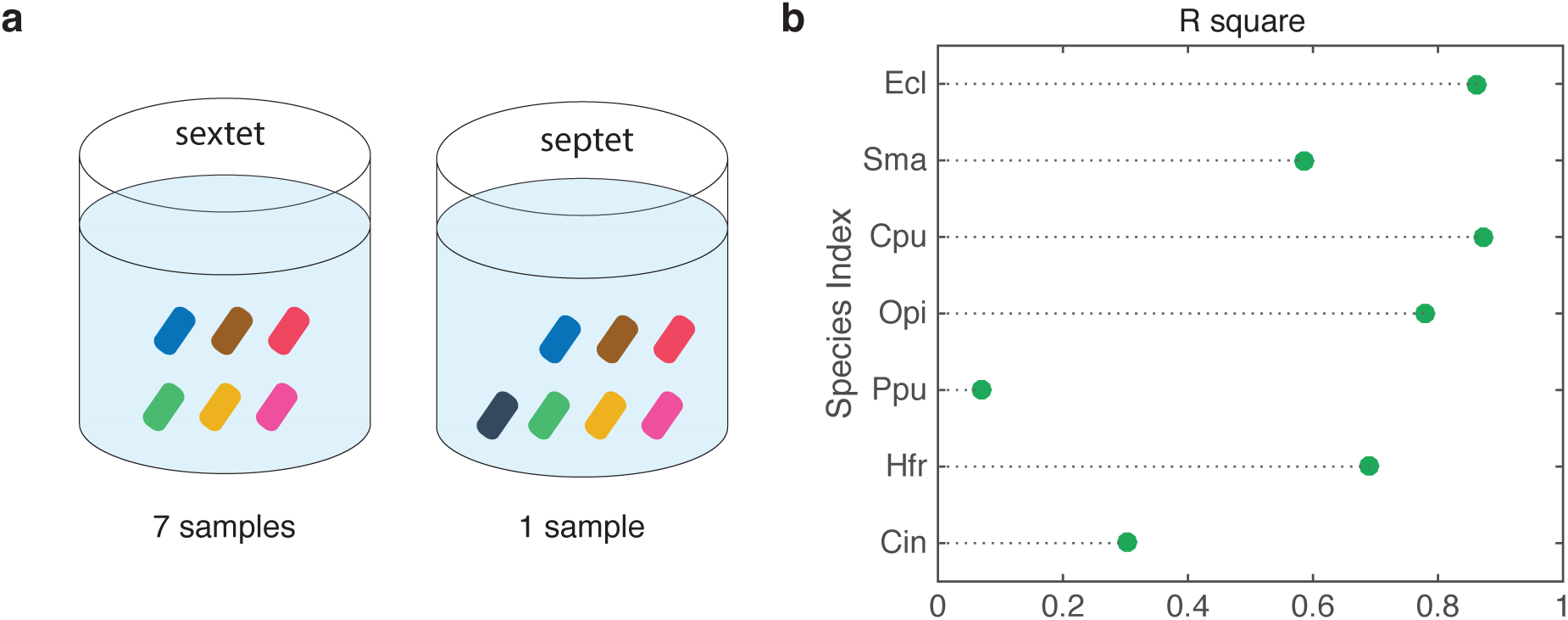
Inferring interaction types in a synthetic community of maize roots with 7 bacterial species. **a.** 8 different species combinations: all 7 sextets and 1 septet. **b.** We find that *R*^2^ of all fitted hyperplanes are smaller than 0.9. This indicates that the given samples cannot be properly described by the GLV model, and we should focus on the inference of interaction types.

**SUPPLEMENTARY FIGURE 12.**
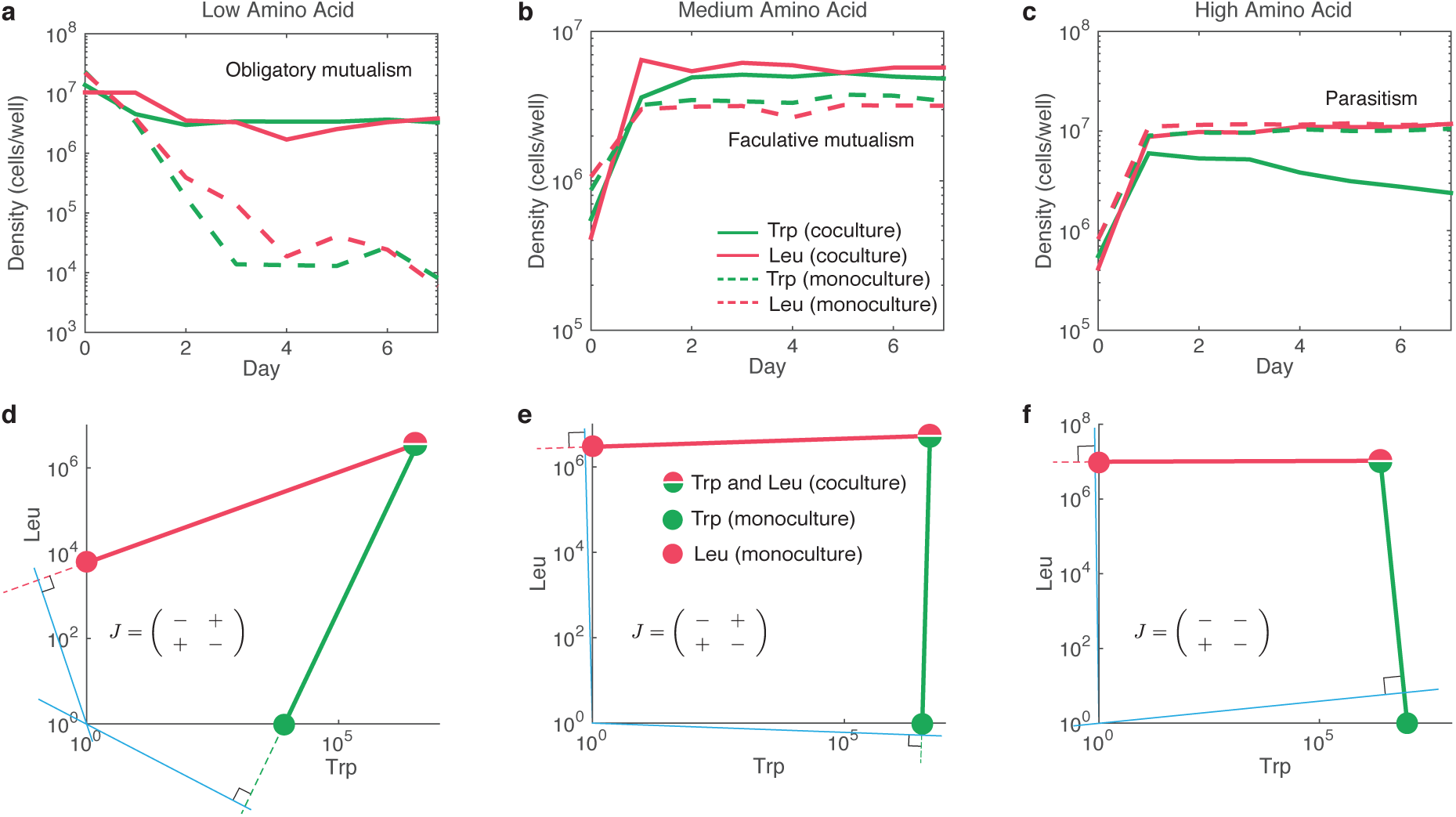
Inferring interaction types in a synthetic microbial community of two cross-feeding partners with different amount of resource availability. **a-c.** The abundance of the co-cultures (solid line) and monocultures (dashed line) for the Trp (green) and Leu (red) strains with the resources of low, medium and high amino acid. Leu and Trp strains are auxotrophic for each other. However, their co-cultures under different amount of resources exhibit different interaction types. **d-f.** Diagrams of our inference method. The inferred results are consistent with the experimental observations.

**SUPPLEMENTARY FIGURE 13.**
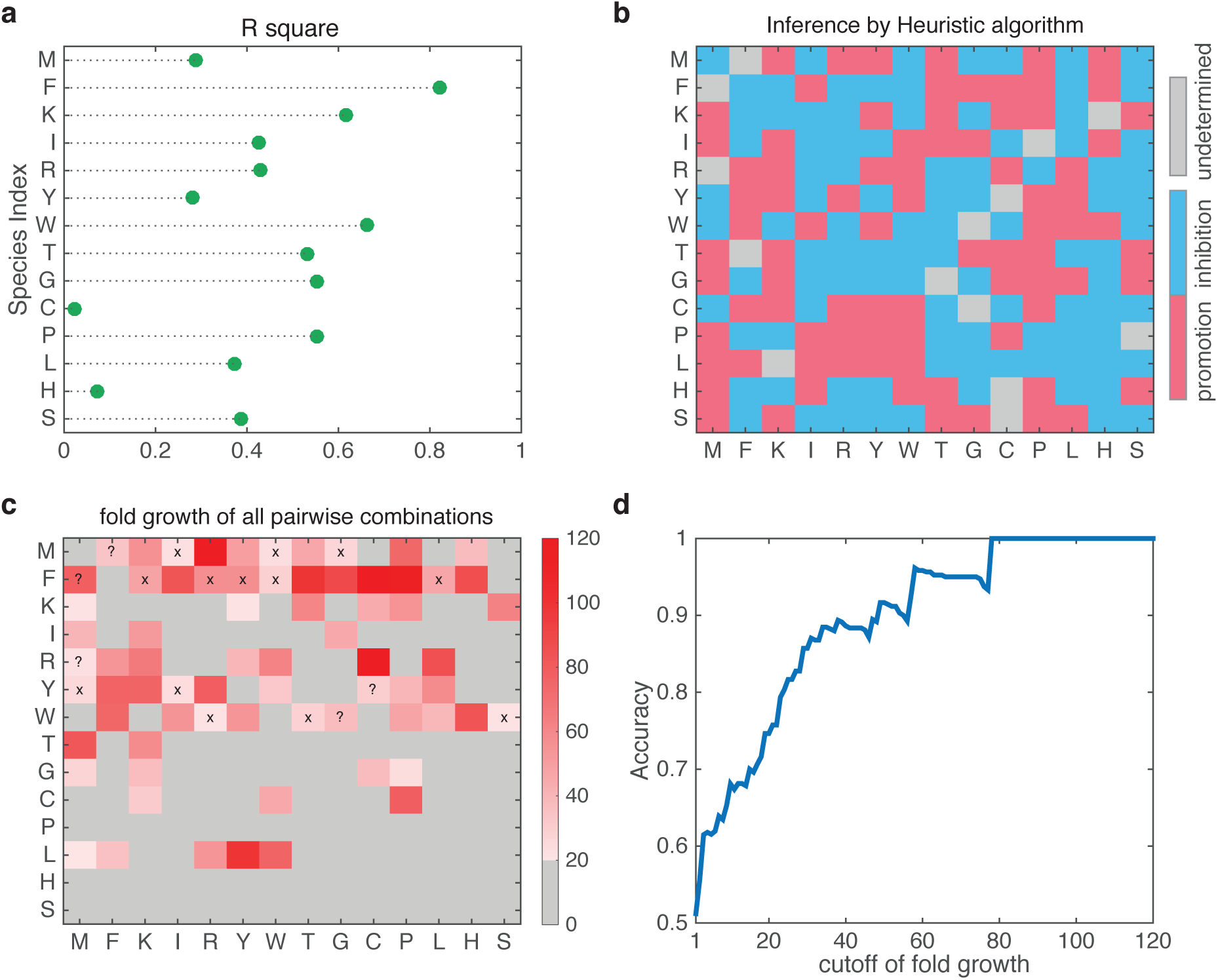
Inferring interaction types in a synthetic community of 14 strains each containing a gene knockout that lead to an auxotrophic phenotype of 1 of 14 essential amino acids. The dataset consists of 91 steady-state samples, each involving a particular pair of the 14 strains. **a.** We find that *R*^2^ of all fitted hyperplanes are smaller than 0.9, suggesting that this community cannot be properly described by the GLV model. **b.** Inferred interaction types by our heuristic algorithm (with 1000 user-defined intersection lines) using only 91 steady-state samples. **c.** The experimentally measured fold growth matrix *F* = (*F*_*ij*_), with *F*_*ij*_ the fold growth of strain *i* (row) in the co-culture paired with strain *j* (column). We set *F*_*ij*_ ≥ 20 as the indication of promotion effect of strain *j* on strain *i*. There are in total 71 promotion interactions with such a large confidence (shown in red). Among them, 53 were correctly inferred, 13 (marked as ‘×’) were not correctly inferred, and 5 (marked as ‘?’) were undetermined by our method, resulting in accuracy 53/71 = 74.65%, at this particular fold growth threshold. **d.** The inference accuracy of promotion effect as a wide range of threshold value of fold growth.

## Footnotes

1 We say a steady state is feasible if it belongs to the orthant 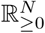, that is, if no taxon has negative abundance.

